# Incorporating Target-Specific Pharmacophoric Information Into Deep Generative Models For Fragment Elaboration

**DOI:** 10.1101/2021.10.21.465268

**Authors:** Thomas E. Hadfield, Fergus Imrie, Andy Merritt, Kristian Birchall, Charlotte M. Deane

## Abstract

Despite recent interest in deep generative models for scaffold elaboration, their applicability to fragment-to-lead campaigns has so far been limited. This is primarily due to their inability to account for local protein structure or a user’s design hypothesis. We propose a novel method for fragment elaboration, STRIFE that overcomes these issues. STRIFE takes as input Fragment Hotspot Maps (FHMs) extracted from a protein target, and processes them to provide meaningful and interpretable structural information to its generative model, which in turn is able to rapidly generate elaborations with complementary pharmacophores to the protein. In a large-scale evaluation, STRIFE outperforms existing, structure-unaware, fragment elaboration methods in proposing highly ligand efficient elaborations. In addition to automatically extracting pharmacophoric information from a protein target’s FHM, STRIFE optionally allows the user to specify their own design hypotheses.

## Introduction

Fragment-based drug discovery (FBDD) approaches are increasingly being used for the rational design of novel compounds.^1,2^ FBDD campaigns aim to identify smaller-than-drug-like molecules which bind weakly to the target and use them as a basis for developing a high-affinity binder. Compared to traditional design methodologies, FBDD methods have a number of advantages. First, starting from small fragments with low molecular weight allows a greater degree of control over the physical properties of the resulting molecule than using a drug-like small molecule as a starting point.^3^ They also facilitate a more efficient exploration of chemical space, with a 2005 study^4^ reporting that hit rates for fragment libraries were 10-1000 times higher than standard high-throughput screening assays. Fragment-based approaches therefore offer a higher chance of identifying a starting point and enhanced control over the subsequent optimisation process.

Following the identification of a set of fragment hits against a target from a fragment screen, there are three main strategies for developing a lead molecule with high binding affinity:^5^ The first, elaboration (or growing), involves selecting a single fragment and adding functional groups to form further favourable interactions with the protein. The second, fragment linking, takes two fragments bound concurrently in the same region of the protein and designs a molecular bridge between them such that the resulting molecule contains both fragments as substructures. Finally, fragment merging requires two or more fragments to bind in overlapping regions and involves the design of molecules which incorporate motifs from each fragment.

In each case, designs are currently proposed on an ad-hoc basis by human experts who draw on standard computational techniques and their deep understanding of chemistry to generate promising ideas. However, human experts may be hindered by implicit biases from past successes and failures, and when working with a large number of hits from a large fragment screen it will not be feasible for a human expert to objectively assess all possible elaboration opportunities for suitability.

Several authors have recently proposed deep-learning based methods to help improve the efficiency of fragment-to-lead campaigns. Graph-based approaches for scaffold elaboration were proposed by Lim et al. ^6^ and Li et al. ^7^, which provide a model with a fragment and generate a set of molecules which contain the original fragment as a substructure, whilst Arús-Pous et al. ^8^ proposed a SMILES-based^9^ model, Scaffold-Decorator, which gave the user the ability to decide which atoms in the fragment should be used as an exit vector, allowing greater control over the types of elaborations generated. However, none of the above approaches allow for the specification of a preferred elaboration size, which, combined with their inability to account for protein structure when generating elaborations, means they cannot ensure that elaborations made by the model would be of an appropriate size to fit within the binding pocket. More recently, we proposed DEVELOP,^10^ a fragment-based generative model for linking and growing which built on our DeLinker^11^ model. DEVELOP allows the specification of pharmacophoric constraints and linker/elaboration length, providing a greater degree of control over the resulting molecules. In concurrent work to DEVELOP, Fialková et al. ^12^ proposed LibINVENT, an extension to Scaffold-Decorator^8^ which can be used to design core-sharing chemical libraries using only specific chemical reactions. LibINVENT also allows users to generate molecules with high 3D similarity to an existing active molecule via reinforcement learning. However, both DEVELOP and LibINVENT are reliant on either a pre-existing active or human specification of pharmacophoric constraints to generate targeted sets of molecules, making them more suitable tools for R-group optimisation than for designing compounds against a novel target.

Orthogonal to the generative approaches described above, several recent papers have proposed database-based approaches to compound design. A recent method, CReM,^13^ is based on the idea that a fragment within the context of a larger molecule can be interchanged with another fragment that has been observed to have the same local context in another molecule. CReM identifies potential elaborations by searching a database of molecules for fragments which have the same local context as the specified exit vector. Other recent database-based approaches incorporate protein-specific information: FragRep^14^ takes a protein and ligand as input and enumerates modifications to the ligand by cutting the ligand into fragments and replaces a fragment with similar fragments from a database which would preserve the same protein-ligand interactions, whilst DeepFrag^15^ uses a structure-aware convolutional neural network to select the most appropriate elaborations from a database of possible elaborations.

For the task of generating molecules ‘from scratch’, a number of authors have proposed generative models which extract information directly from the protein. Skalic et al. ^16^ used a GAN^17^ to generate ligand shapes complementary to the binding pocket which were then used to generate potential molecules by employing a shape-captioning network. Masuda et al. ^18^ encoded atomic density grids into separate latent representations for ligand and protein and trained a model to generate 3D ligand densities conditional on the protein structure, which were then translated into discrete molecular structures. Whilst both papers demonstrated that the ligands generated by their respective models were dependent on the learned structural representations, the models do not facilitate the specification of a design hypothesis provided by a human expert. Kim et al. ^19^ used water pharmacophore models to learn the location of key protein pharmacophores which were then used to construct a training set of molecules with complementary pharmacophores. Whilst this approach would more readily integrate into standard drug-discovery efforts, it requires the training of a separate deep learning model for every target, as each target requires a training set of compounds which match the water pharmacophores.

In this work we propose STRIFE (**Str**ucture **I**nformed **F**ragment **E**laboration), a generative model for fragment elaboration which extracts interpretable and meaningful structural information from the protein and uses it to make elaborations. This is different to all existing fragment-based generative approaches which either extract information implicitly from known ligands or do not make use of any protein-specific information when generating molecules. To allow straightforward integration into fragment-to-lead campaigns, STRIFE is readily customisable; in addition to the design hypotheses extracted directly from the protein, we provide a simple-to-use functionality which allows users to specify their own design hypotheses and generate elaborations with the aim of satisfying a desired pharmacophore. In a large-scale evaluation derived from the CASF-2016 set,^20^ we show that STRIFE offers substantial improvements over existing fragment-based models.^8,13^ We further demonstrate the applicability of STRIFE to real-world FBDD campaigns through two fragment elaboration tasks derived from the literature. In the first, we make elaborations to a fragment bound to N-myristoyltransferase, a key component in rhinovirus assembly and infectivity, and show that STRIFE is able to generate several elaborations that are strikingly similar to a highly potent inhibitor.^21^ To demonstrate how user-specified design hypotheses can be incorporated into STRIFE, we consider the fragment-inspired small molecule inhibitor of tumour necrosis factor reported by O’Connell et al. ^22^. In this example, the elaboration proposed by O’Connell et al. ^22^ induces a substantial movement in a Tyrosine side chain. We manually specified a design hypothesis to explore side-chain flexibility, and successfully recovered the elaboration proposed by O’Connell et al. ^22^, as well as a range of other elaborations which were predicted to induce a similar movement in the Tyrosine side chain.

## Methods

We present our deep generative model for fragment elaboration, STRIFE, which requires the user to specify a target protein, a bound fragment, and the fragment exit vector. In our previous work,^10^ we demonstrated how the imposition of pharmacophoric constraints allowed a substantial degree of control over the types of functional groups added to a fragment. STRIFE builds on the approach proposed in Imrie et al. ^10^, where the pharmacophoric constraints were extracted from existing active molecules, by extracting pharmacophoric constraints directly from the protein, thereby extending its applicability to a much broader range of targets. Pharmacophoric information is extracted by calculating a Fragment Hotspot Map^23^ (FHM), which describes regions of the binding pocket that are likely to make positive contribution to binding affinity. STRIFE then identifies pharmacophoric constraints which are likely to place a pharmacophore within a matching hotspot region and uses the pharmacophoric constraints to generate elaborations.

### Fragment Hotspot Maps

We calculate FHMs using the Hotspots API^24^ which implements the algorithm described by Radoux et al. ^23^; in this work all FHMs were calculated using the default parameters given by Curran et al. ^24^. An FHM is calculated as follows: Atomic Propensity Maps are calculated using SuperStar, ^25^ which defines a grid covering the protein with equally spaced points 0.5Å apart, and uses data from the Cambridge Structural Database (CSD)^26^ to assign a propensity for a given probe type at each grid point; If an interaction between two groups at a certain distance and angle is particularly favourable then it will occur more frequently in structures stored in the CSD and be assigned a higher propensity score. Once an Atomic Propensity Map has been calculated an FHM is derived by first weighting the scores assigned to each grid point in proportion to how buried in the protein the grid point is.

The FHM scores are then calculated by using small chemical probes which take the form of an aromatic ring with different atoms in the substituent position; for the apolar hotspot maps the substituent is a methyl group, whilst for the acceptor and donor hotspot maps the substituent is a carbonyl and amine, respectively. The probes are translated to all grid points with weighted propensity scores above 15 and randomly rotated 3000 times about the center of the substituted atom. For each pose, each atom receives a score read from the weighted propensity map and the probe scores are calculated as the geometric mean of the atom scores; as an atom receives a score of zero if it clashes with the protein, the geometric mean gives a score of 0 to any pose which clashes.

FHMs have a number of attractive properties. As only grid points with an above-threshold weighted propensity score are sampled, and the propensity scores are weighted by how buried in the protein they are, regions of the protein which are overly exposed are unlikely to be identified as hotspot regions. Additionally, because probe poses which clash with the protein attain a score of zero, any region identified as a fragment hotspot must be able to accommodate a molecule of reasonable size, meaning that the risk of attempting to satisfy a pharmacophore identified by the FHM which cannot be accessed by an elaboration is reduced.

### FHM Processing

For a protein target, STRIFE uses FHMs to guide the generative model in the placement of functional groups which can interact with the target. As the different hotspot maps are used for different purposes, they are processed slightly differently (Figure 1): the acceptor and donor hotspots are used to identify desirable pharmacophoric constraints, whilst the apolar maps are used to verify that the fragment is located in an appropriate binding site. For the apolar maps, we identify all grid points which have a value greater than 1 and discard all other points. Similarly, for the acceptor and donor maps, we retain all grid points which have a value greater than 10. Whilst Radoux et al. ^23^ reported that values greater than 17 were generally predictive of fragment binding, we selected 10 as a threshold to obtain wider coverage; this parameter is simple to change to restrict the search to higher quality hotspots.

**Figure 1:**
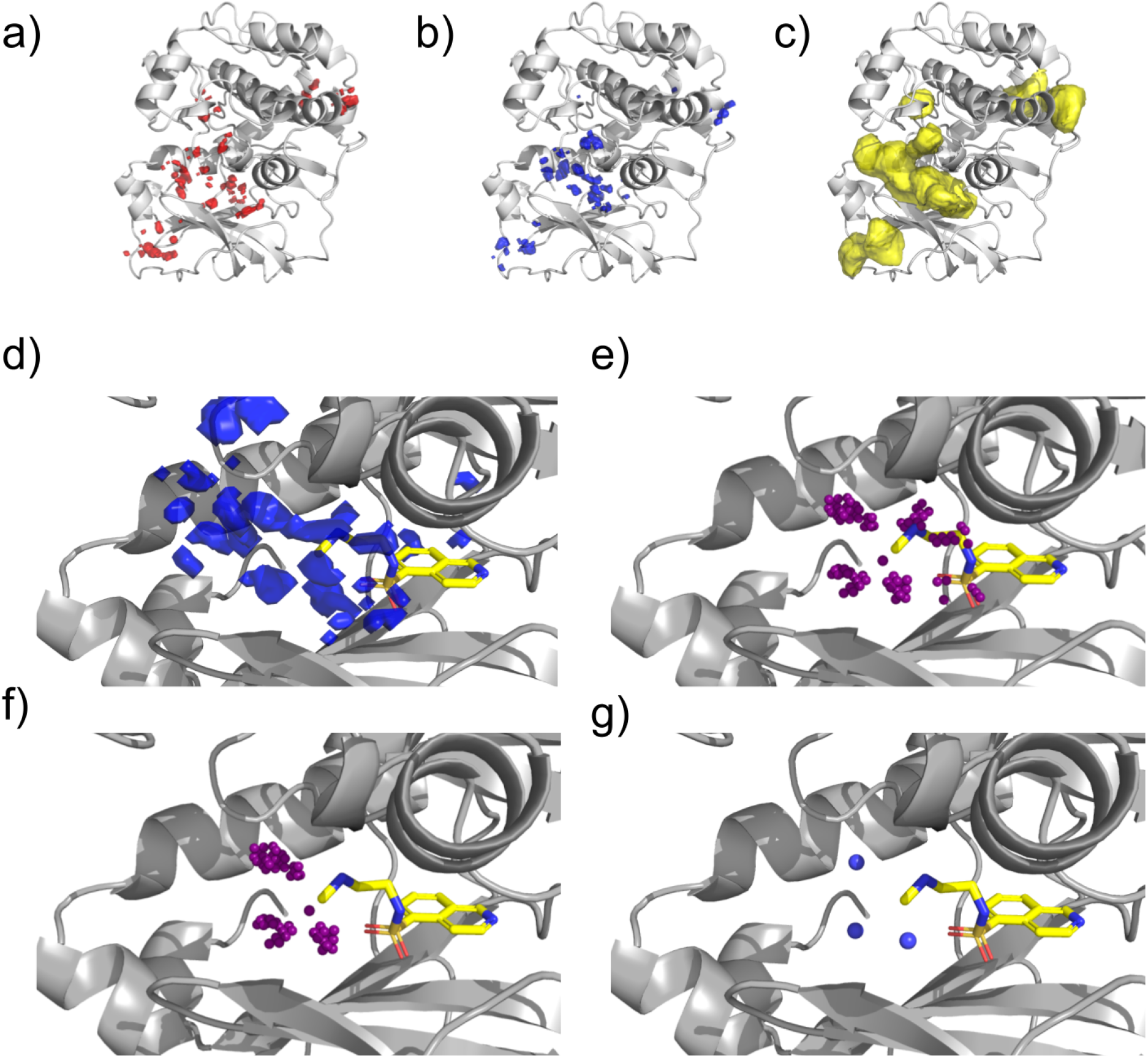
Processing Fragment Hotspot Maps. a) Acceptor Hotspot Map. b) Donor Hotspot Map. c) Apolar Hotspot Map. A matching pharmacophore placed within a hotspot has a chance of making a disproportionate contribution to binding affinity. d) An unprocessed donor hotspot map in the vicinity of the fragment of interest. e) Each sphere represents a voxel in the hotspot map. Voxels which are too far away from the fragment exit vector are discarded. f) Voxels which are closer to another fragment atom than the exit vector are removed. g) Voxels are clustered based on their position. STRIFE attempts to generate elaborations such that a matching ligand pharmacophore is in close proximity to a cluster centroid.

To process the acceptor and donor maps, all points which are less than 1.5Å or greater than 5Å from the fragment exit vector are discarded, to allow for elaborations of appropriate length; these distance thresholds were chosen to reflect the iterative nature of a fragment-to-lead campaign, where practitioners typically make a succession of small elaborations, but they can be altered by the user to admit longer or shorter elaborations.

A greedy clustering algorithm is employed to identify contiguous hotspot regions as follows: A cluster is initialised as a single point and all unclustered points which are within 1Å of the grid point are added to the cluster. For each point in the cluster, the distance to all remaining unclustered points is calculated and any points which are within 1Å are added to the cluster until no unclustered points can be added. Once a cluster has terminated, a new cluster is defined by selecting a single unclustered point, until all points have been assigned to a cluster. For each hotspot cluster, centroids are defined by computing the mean position of the points in the cluster. To reduce redundancy, if two cluster centroids are closer than 1.5Å apart the cluster centroid corresponding to the smaller cluster is deleted. In addition, if a cluster is smaller than eight points it is removed, unless no clusters of eight or more points exist, in which case smaller clusters are retained.

We use the apolar maps to conduct a final filtering step, adopting the heuristic that a molecule which is entirely contained within an apolar hotspot region has a better chance of binding to the protein. Therefore, if an acceptor or donor cluster centroid is not contained within a hotspot region then it is filtered out. Additionally, if all fragment atoms are not contained within an apolar hotspot then we consider the fragment to be unsuitable for elaboration and terminate the algorithm. Whilst this might appear to be overly restrictive, in practice the apolar hotspot maps typically cover the majority of binding sites in a target and this filtering step can be easily negated if the user believes that a fragment is a suitable candidate for elaboration.

The final output of the processing scheme are the 3D coordinates of the remaining cluster centroids from the acceptor and donor maps (here-after “pharmacophoric points”). In the subsequent molecule generation steps, our aim is to generate elaborations which place matching functional groups in close proximity to the pharmacophoric points. Whilst the above pipeline automates the process of defining pharmacophoric points, we also provide a simple-touse functionality for users to define their own pharmacophoric points, allowing them to pursue a range of different design hypotheses (see Methods, Customisability).

Next, we describe how STRIFE uses a set of pharmacophoric points to generate elaborations with complementary pharmacophores to the target.

### Generative Model

The generative model employed by STRIFE is similar to our previous work, DEVELOP,^10^ where the generative process is based upon the Constrained Graph Variational Autoencoder framework proposed by Liu et al. ^27^. STRIFE differs from DEVELOP in the structural information ***D*** provided to the model when decoding molecules (see Methods, STRIFE Algorithm). Given a fragment, ***f***, and structural information, ***D***, elaborations are generated as follows: Representing ***f*** as a graph, each node *v* is assigned an *h*−dimensional vector representation **z**_*v*_ and corresponding label *l_v_*, denoting the atom type of the node. A set of *K* ‘expansion nodes’, 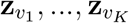 are generated by sampling from an *h* − dimensional standard normal distribution and each expansion node is assigned a label 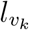 by a linear classifier which takes 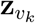 and ***D*** as input. The expansion nodes represent the possible atoms which may be appended to the fragment.

Starting from the fragment exit vector, the model samples a node to add to the graph from the set of expansion nodes. To choose whether to form a bond between node *v* and node *u*, we use a neural network which takes as input:

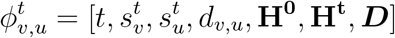

Where:

- *t* is the number of time steps that have currently been taken.
- 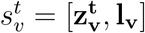 is the concatenation of latent vector and label at the *t^th^* time step.
- *d_u,v_* is the graph distance between *u, v*
- **H^j^** is the average of all latent vectors at the *j^th^* time step.

After a new node has been added to the graph, a gated graph neural network^28^ is used to update the encodings for each node, to reflect its potentially altered neighbourhood. This iterative approach continues until termination where the final molecule is returned. For additional details regarding the generative model framework, can be found in our previous work.^10,11^

### STRIFE Algorithm

Above we described how STRIFE uses Fragment Hotspot Maps (FHMs)^23^ to obtain an interpretable representation of structural information and how, given a fragment, *f*, and structural information, *D*, we can generate elaborations to the fragment. Here we describe how these processes fit within the STRIFE algorithm. In particular, we outline how the 3D pharmacophoric points derived from the FHMs are converted to a representation of structural information, *D*, which is used to generate elaborations.

The structural information ***D*** can be provided to the generative model in two different forms: The first is a coarse-grained pharmacophoric representation, where the model is simply provided with a vector containing the number of Hydrogen Bond Acceptors, the number of Hydrogen Bond Donors, and the number of Aromatic groups. The desired pharmacophoric profile of the generated elaborations can also be more precisely specified by adding the predicted path distances (the length of the shortest sequence of atoms connecting two points) from the exit vector to the pharmacophore, providing a greater degree of control over the types of elaborations made by the model. STRIFE utilises both of these representations of pharmacophoric information at different stages of the algorithm: In the Exploration phase (Figure 2a), STRIFE uses the coarse-grained representation to generate a wide range of elaborations, which are then assessed for suitability. In the Refinement phase (Figure 2b), fine-grained pharmacophoric profiles are derived from the most suitable elaborations and are used to generate further elaborations. Additional details are provided below and in the Supplementary Information.

**Figure 2:**
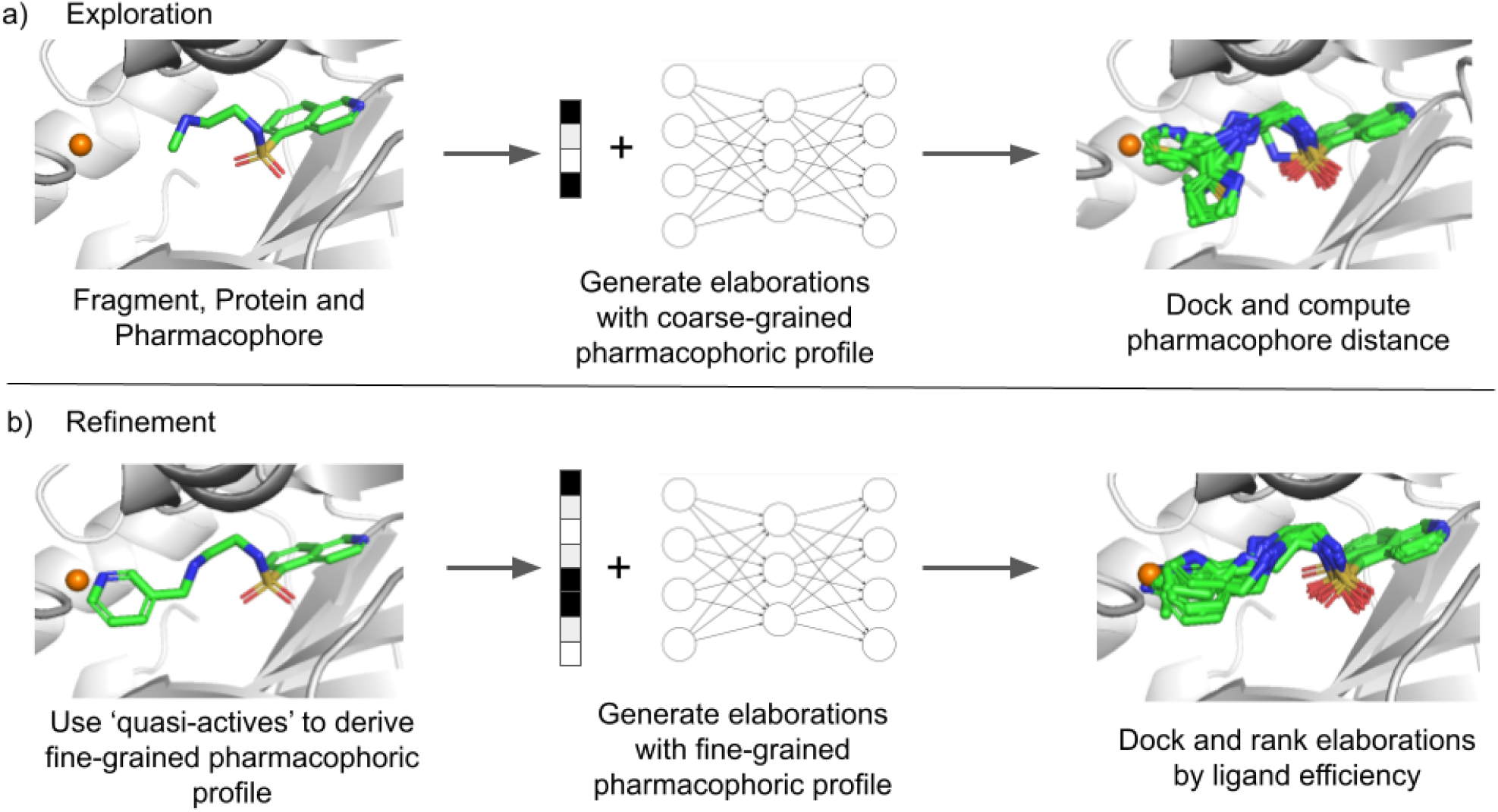
Illustration of how STRIFE generates elaborations which place pharmacophores close to a specified pharmacophoric point. a) STRIFE first generates elaborations using a coarse-grained pharmacophoric profile and docks them using the constrained docking functionality in GOLD. ^30^ b) Elaborations which placed a matching pharmacophore in close proximity to the pharmacophoric point are used to derive a fine-grained pharmacophoric profile. STRIFE then generates elaborations using those pharmacophoric profiles; the resulting molecules are docked and ranked by their predicted ligand efficiency.

In a standard fragment elaboration campaign, where practitioners typically work in an iterative way, making small elaborations to a fragment which is then optimised before making additional elaborations to the optimised molecule. In this paper we demonstrate STRIFE generating elaborations which place a pharmacophore close to a single pharmacophoric point at a time. For example, if the set of pharmacophoric points contains one donor and one acceptor, STRIFE will attempt to produce a set of elaborations which include a donor in close proximity to the donor pharmacophoric point and a set of elaborations which place an acceptor in close proximity to the acceptor pharmacophoric point, but will not attempt to satisfy both pharmacophoric points simultaneously. STRIFE is capable of attempting to satisfy multiple pharmacophoric points simultaneously, but this is not recommended unless the pharmacophoric points have been manually specified or inspected by the user, as it may not be possible to simultaneously satisfy certain combinations of pharmacophores with a single elaboration. After obtaining a series of pharmacophoric points from the FHM, STRIFE proceeds as follows:

#### Exploration Phase

STRIFE aims to generate a set of elaborations which contain functional groups in close proximity to a pharmacophoric point. To facilitate this, for each pharmacophoric point we predict the atom-length distance between the fragment exit vector and the pharmacophoric point using a trained support vector machine.^29^ The atom-length prediction is then used to control the length of elaborations proposed by STRIFE, whilst the pharmacophoric profile of the generated molecules is controlled by the coarse-grained representation representation described above. As the coarse-grained pharmacophoric profile doesn’t specify a desired path distance between the exit vector and the ligand pharmacophore, we obtain a broad range of different elaborations. The elaborations are filtered (see Supplementary Information) and docked using the constrained docking functionality in GOLD, ^30^ where the structure of the fragment is provided as the constraint. Each molecule is docked 10 times and the top-ranked pose selected. For each top-ranked pose, we compute the distance between the 3D pharmacophoric point and a matching pharmacophore in the molecule. We then identify all molecules where the resulting distance is less than 1.5Å and select the five molecules for which the distance between pharmacophoric point and ligand pharmacophore is smallest. If less than five molecules exhibit a distance of less than 1.5Å, we select only molecules which fulfil this criteria.

#### Refinement Phase

The molecules which exhibit a functional group in close proximity to a pharmacophoric point provide useful information, as they can be used to derive the more fine-grained representation of structural information which specifies the path distance between the exit vector and each ligand pharmacophore; as such, we refer to these molecules as “quasi-actives”, because they play a similar role to known actives in existing generative models. Having obtained a set of quasi-actives for each pharmacophoric point and used them to derive a set of structural information vectors ***D***_1_*, **D***_2_*, …, **D**_n_*, the user can either generate a fixed number of elaborations using each ***D**_i_* or request a fixed total number of elaborations, where a structural information vector is randomly sampled from 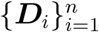 for each elaboration. As before, the generated molecules are filtered and docked using the constrained docking functionality in GOLD. ^30^ Finally, each unique molecule, *m*, is ranked by its ligand efficiency, computed as the docking score divided by the number of heavy atoms, allowing the user to quickly prioritise a small number of elaborations for consideration.

### Customisability

Although STRIFE can automatically extract a set of pharmacophoric points from a protein, in a real-world drug discovery setting practitioners may wish to explore their own design hypotheses. To facilitate such usage, we provide a simple-to-use functionality which allows a user to manually specify the location of a pharmacophore in the context of the protein. The tool, shown in Figure 3, loads a lattice centered around the fragment exit vector into a molecule viewer: To manually specify their own pharmacophoric profiles, the user simply selects the lattice points corresponding to their desired pharmacophore location, saves the resulting object and runs STRIFE as usual.

**Figure 3:**
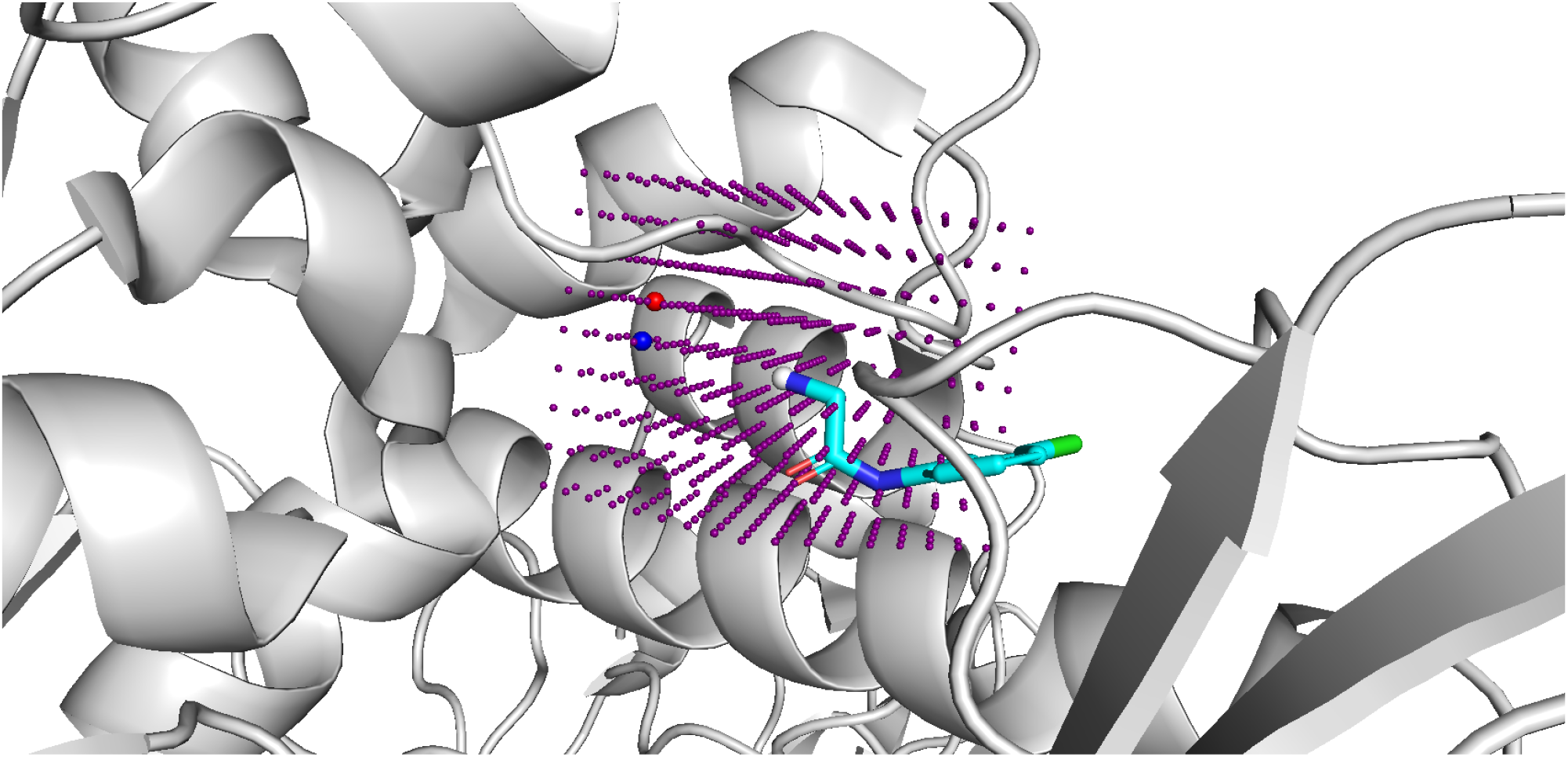
Example of how the pharmacophoric points provided to STRIFE can be customised using the molecule viewer PyMOL.^31^ A lattice of points are centered about the fragment exit vector (denoted by the grey atom), and the user simply selects the point(s) they wish to denote as a pharmacophoric point and saves them in an SDF file. The red and blue points represent an Acceptor point and Donor point, respectively. STRIFE can then be run as usual and will attempt to make elaborations which places matching pharmacophores close to the user-specified pharmacophoric points.

### Model Training

We trained our generative models using a training set derived from the subset of ZINC^32^ randomly selected by Gómez-Bombarelli et al. ^33^. For each molecule, we obtained a series of fragment-ligand pairs by enumerating all cuts of acyclic single bonds which were not part of functional groups. The resulting training set comprised approximately 427,000 examples. The same hyperparameters were used for training as in our previous work.^10^

### Experiments

We assessed the ability of our model to make appropriate elaborations using a test set derived from the CASF-2016 set.^20^ This test set was constructed using the same procedure used to generate our training set and initially comprised 237 examples. As our aim was to assess the ability of our model to learn from the structural information supplied by the FHMs, we excluded from our test sets examples where the ground-truth molecule was not contained within an apolar hotspot region and examples where no suitable pharmacophoric points could be identified by the Hotspots algorithm. In addition, we filtered examples where STRIFE was unable to identify any quasi-actives. These filtering steps removed 109, 26 and 1 examples from the test set respectively, leaving a final test set of 101 examples (a full list is given in Table S1). Whilst the filtering steps outlined above removed a substantial proportion of examples from our test set, the initial test set was constructed by fragmenting the ground truth ligand without consideration of the associated protein. As such, many of the fragments would not have been considered suitable candidates for elaboration.

Using the STRIFE pipeline (Figure 2), we sampled a set of 250 elaborations for each example in the test set. We compared STRIFE to three baselines: The model published by Arús-Pous et al. ^8^, “Scaffold-Decorator”, the database-based CReM, and a truncated version of the STRIFE algorithm (STRIFE_*NR*_) which generated elaborations from the coarse-grained model (essentially only conducting the Exploration phase from Figure 2a and omitting the Refinement phase). We provided CReM with the same set of 250k molecules we used to derive the training sets for STRIFE, which was converted into a database of fragments using CReM’s fragmentation procedure. The Scaffold-Decorator model was trained using the same set of examples as the STRIFE generative models.

### Evaluation Metrics

For our experiments on the CASF test set, we report several standard 2D metrics in line with those reported in our previous work:^10^

- **Validity**: Proportion of generated molecules which could be parsed by RD-Kit^34^ and for which at least one atom was added to the fragment.
- **Uniqueness**: The proportion of distinct molecules generated by the model, calculated as the number of distinct molecules divided by the total number of molecules.
- **Novelty**: The proportion of generated molecules for which the elaboration was not included in the model training set.
- **Passed 2D Filters**: The proportion of generated molecules which passed a set of 2D filters. A generated molecule was filtered out if the SAScore^35^ of the generated molecule was higher (harder to synthesise) than the SAScore associated with the fragment, if the elaboration contained a non-aromatic ring with a double bond or if the molecule failed to pass any of the pan-assay interference (PAINS)^36^ filters.

We did not compute the proportion of unique or novel associations proposed by CReM, as CReM does not allow the specification of a desired number of elaborations: CReM returns the set of elaborations contained in the database deemed ‘reasonable’, meaning that all elaborations proposed by CReM are by design unique. Similarly, as CReM proposes molecules from a fixed vocabulary of possible elaborations, none of the elaborations proposed by CReM could be considered novel.

To assess the ability of STRIFE to generate elaborations capable of forming promising interactions with the target, we used the constrained docking functionality in GOLD ^30^ to dock each generated ligand 10 times and calculated the docking score of the top-ranked pose for each ligand. To mitigate the tendency of classical scoring functions to favour larger molecules over smaller ones,^37^ we calculated the ligand efficiency of each molecule by dividing the docking score by the number of heavy atoms. To account for the variation in docking scores across different targets, we standardise the ligand efficiencies attained by a model on a specific example to have zero mean and unit variance, applying the same transformation to the ground truth ligand efficiency. For the *j^th^* example, we compute ΔSLE_*α,j*_ = SLE_*α,j*_ − SLE_*GT,j*_, where SLE_*α,j*_ is the average standardised ligand efficiency of the top *α* ranked molecules and SLE_*GT,j*_ is the standardised ligand efficiency of the corresponding ground truth. If *α* is specified as greater than the total number of elaborations for which the ligand efficiency was computed (as only molecules which pass the 2D filters are docked), we use the average standardised ligand efficiency of all such molecules. We average over all examples to obtain 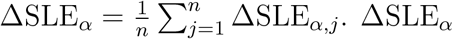 (Standardised Ligand Efficiency Improvement) only considers a subset of the molecules generated by each model, mirroring how a large number of molecules produced by a generative model would be assessed in a real-world fragment-to-lead campaign, where it is unlikely that a human expert would manually inspect hundreds of lowly ranked molecules.

As CReM is unable to return a fixed number of elaborations, we calculated three sets of summary statistics for CReM, each using a different subset of the test set. In all cases, if CReM returned more than 250 elaborations for a specific example, we sampled a set of 250 elaborations from the larger set:

- The set of examples for which CReM returned 250 elaborations (*n* = 45).
- The set of examples for which CReM returned 50 or more elaborations (*n* = 62).
- The set of examples for which CReM returned at least one elaboration (*n* = 82).

We present the results for the first set in Table 1 and compare the results between the three subsets in the Supplementary Information (Table S2): in the case where we included all examples with at least one elaboration, the ΔSLE_*α*_ values were substantially degraded by the subset of examples where only a small number of elaborations were proposed.

**Table 1:** Comparison of CReM, Scaffold-Decorator, STRIFE_*NR*_ and STRIFE on the CASF test set (see Methods, Evaluation Metrics for definitions of the metrics). **Bold** indicates the best value obtained across the different methods.

| Metric | CReM | Scaffold-Decorator | STRIFE<sub>NR</sub> | STRIFE |
| --- | --- | --- | --- | --- |
| Valid | **100%** | 99.98% | 99.5% | 98.96% |
| Unique | N/A | 32.78% | **56.96%** | 37.31% |
| Novel | N/A | 4.23% | **55.65%** | 49.21% |
| Pass 2D filters | 66.06% | **98.2%** | 73.81% | 75.38% |
| $\Delta\text{SLE}_{20}$ | -0.029 | 0.1 | 0.44 | **0.512** |
| $\Delta\text{SLE}_{50}$ | -0.489 | -0.222 | 0.078 | **0.164** |
| $\Delta\text{SLE}_{100}$ | -0.992 | -0.572 | -0.316 | **-0.228** |

## Results and Discussion

We assessed the ability of STRIFE to propose elaborations to fragments by incorporating meaningful pharmacophoric information into the generative process. Through a large scale evaluation on a test set derived from the CASF-2016 set,^20^ we show that STRIFE is able to generate a wide range of chemically valid elaborations, many of which were not contained in the training set. In addition, in terms of generating elaborations which exhibit high ligand efficiency, STRIFE substantially outperforms existing computational methods for fragment elaboration, ^8,13^ illustrating the advantages of incorporating structural information into the generative model. We demonstrate the applicability of STRIFE to real-world fragment-to-lead campaigns using two case studies derived from the literature; in particular, we show how STRIFE can be used to explore design hypotheses including side-chain movement.

### Large Scale Experiments

Our experiments on the CASF set demonstrate the benefits of including structural information in the generative process (Table 1). All methods generated chemically valid elaborations in more than 99% of cases, illustrating their ability to apply basic valency rules. Scaffold-Decorator, the SMILES-based, structure-unaware generative model proposed by Arús-Pous et al. 8, generated the smallest proportion of unique molecules (33%). STRIFE_*NR*_, a truncated version of the STRIFE algorithm which terminates before the Exploration phase so doesn’t account for the location of fragment hotspots, generated a greater proportion of unique elaborations (57%) than STRIFE (37%). However this is to be expected as the Refinement phase of the algorithm attempts to sample elaborations from a greatly reduced chemical space compared to the Exploration phase.

Illustrating its ability to generalise beyond the information provided in the training set, almost half (49%) of the elaborations proposed by STRIFE were not contained in the training set. By contrast, only 4% of the elaborations generated by Scaffold-Decorator were novel, suggesting that it relies more heavily on the training set when making elaborations. Almost all of the elaborations proposed by Scaffold-Decorator (98%) passed the set of 2D filters, compared to 75% of elaborations generated by STRIFE and 74% by STRIFE_*NR*_. As nearly all of the elaborations proposed by Scaffold-Decorator were contained in the training set, which itself was filtered to remove undesirable elaborations, the high pass rate of 2D filters is unsurprising.

On ΔSLE, which assesses the ability of models to generate elaborations which are more ligand efficient than the ground truth ligand, models that incorporate structural information proposed more ligand efficient elaborations. When considering the top 20 elaborations, the elaborations generated by CReM (ΔSLE_20_ = −0.029) and Scaffold-Decorator (ΔSLE_20_ = 0.1) were on average less ligand efficient than the ground truth, in contrast to STRIFE_*NR*_ (ΔSLE_20_ = 0.44) and STRIFE (ΔSLE_20_ = 0.512). These results indicate that the fine-grained pharmacophoric profiles extracted during the Refinement phase allow STRIFE to generate more ligand efficient elaborations, as the model more often generates elaborations which place pharmacophores in close proximity to a pharmacophoric point. We observed the same trend when the top 50 and 100 elaborations were considered, although in this case the average ligand efficiency obtained by all models was lower than the ground truth ligand efficiency. We show how ΔSLE_*α*_ varies for different values of *α* in the Supplementary Information (Figure S3).

In terms of the proportion of all generated elaborations which were more ligand efficient than the ground truth, STRIFE achieved the largest number, with 26% of elaborations obtaining a higher ligand efficiency than the ground truth, compared to 22%, 17% and 12% for STRIFE_*NR*_, Scaffold-Decorator and CReM (when considering examples with 250 elaborations) respectively.

### Fragment-Based Design of an N-myristoyltransferase Inhibitor

Rhinovirus is a pathogen which plays a key role in complications arising in a variety of important respiratory diseases, including asthma, chronic obstructive pulmonary disease (COPD)^38^ and cystic fibrosis.^39^ Several studies^40,41^ have reported that the host cell’s N-myristoyltransferase (NMT) supports capsid assembly and infectivity, making NMT a potential antiviral drug target.

Following a fragment screen against NMT from the human malaria parasite *Plasmodium falciparum*,^42^ Mousnier et al. ^21^ identified a fragment-like compound, IMP-72 (Figure 4a), with weak (IC_50_ = 20*μM*) activity against Human NMT1 (HsNMT1). The binding mode of IMP-72 was originally determined in NMT from the malaria parasite *P.vivax* (PvNMT), but as the fragment’s key interactions involved residues which are conserved in human NMTs, it was considered to be a viable starting point for the development of an HsNMT1 inhibitor. The authors noted that IMP-72 bound in a region complementary to a previously identified quinoline inhibitor,^43^ MRT00057965, however closer inspection of the overlaid binding modes precluded a fragment merging strategy. To address this the authors constructed a simplified quinoline fragment, IMP-358, which could recapitulate the same interactions as MRT00057965 (S319 in PvNMT and S405 in HsNMT1) without clashing with IMP-72. Despite exhibiting weak inhibition of HsNMT1 (17% at a concentration of 100*μM*), IMP-358 facilitated a synergistic inhibition alongside IMP-72, with the potency of IMP-72 increasing 300-fold for HsNMT1 in the presence of IMP-358. The authors developed a further compound, IMP-917, derived by replacing IMP-358 with a trimethylpyrazole group which was then linked to IMP-72 with an ether linker. Compared to IMP-72, IMP-917 exhibited a 1500-fold improvement in potency (IC_50_ = 0.013*μM*) and retained the key interactions made by both IMP-72 and IMP-358. Finally, the authors made slight modifications to the core of IMP-917 and used the resulting compound to show that NMT inhibition completely prevents rhinoviral replication without inducing cytotoxicity, thereby identifying a potential drug target.

**Figure 4:**
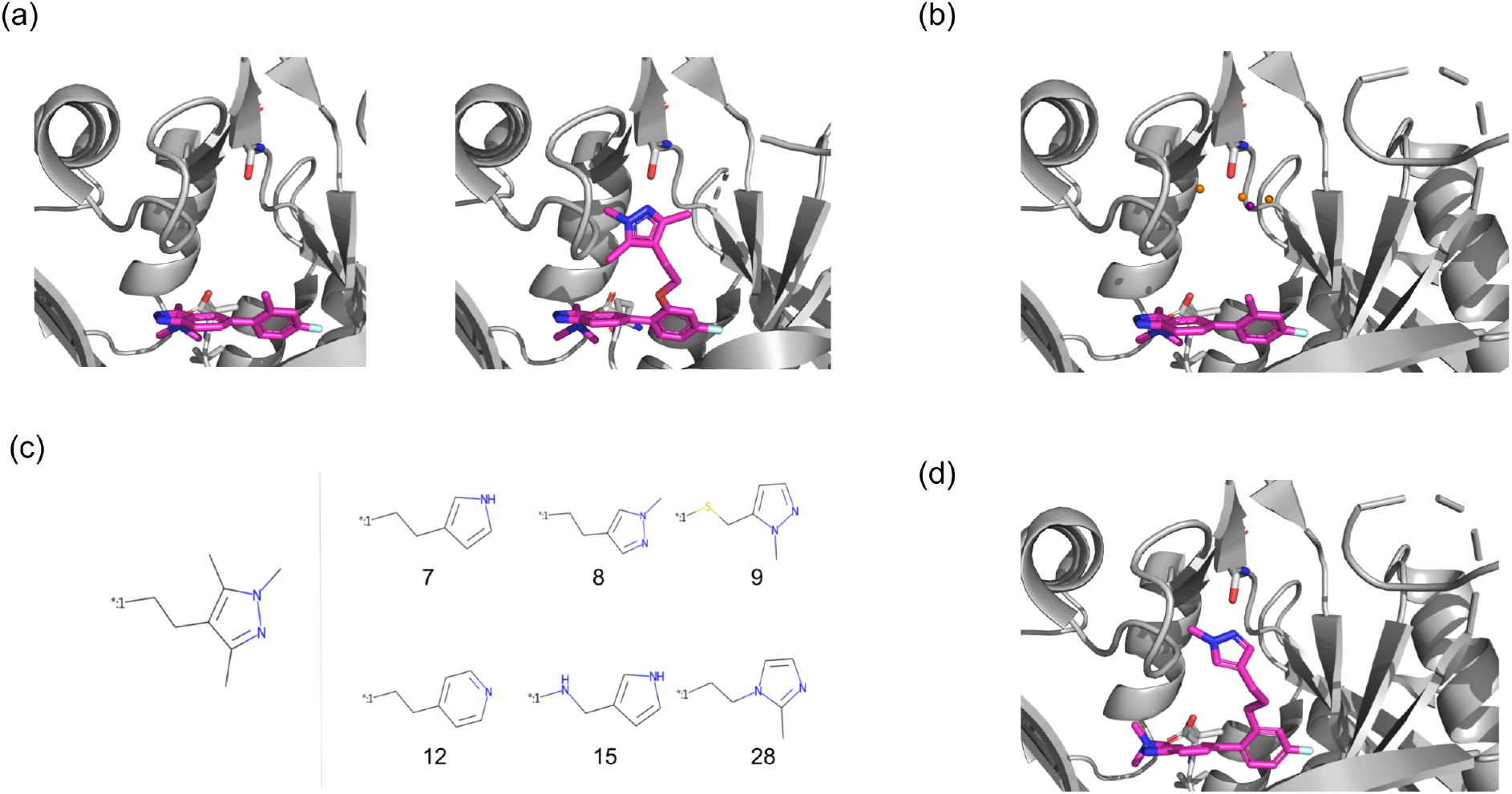
Fragment elaboration case study. (a) Left: Crystal structure (PDB ID 5O48) of the fragment bound to P.vivax NMT. Right: Crystal structure (PDB ID 5O6H) of the optimised compound bound to Human NMT1. The trimethylpyralzole facilitates an interaction with the residue S319. (b) Processed pharmacophoric points from the Fragment Hotspot Map calculated on P.vivax NMT. The orange spheres correspond to hydrogen bond acceptor points whilst the purple sphere corresponds to a hydrogen bond donor point. (c) The elaboration proposed by Mousnier et al. ^21^ (left) compared to several elaborations proposed by STRIFE which satisfied the same design hypothesis (right). The number underneath each elaboration corresponds to the rank assigned to it by STRIFE. (d) Docked pose of one of our elaborations, which appears to be capable of forming the same hydrogen bond interaction with S319.

We investigated the ability of STRIFE to propose molecules that could satisfy the design hypothesis put forward by Mousnier et al. ^21^. Instead of iteratively refining the original quinoline fragment and constructing a linker, we viewed the task as an elaboration problem and sought to propose elaborations which could form interactions with S319. As input to STRIFE, we provided the SMILES string of IMP-72, the exit vector we wished to make elaborations from and the crystal structure of PvNMT (PDB ID 5O48). Although our aim was to design compounds for HsNMT1, we did not have access to a crystal structure of IMP-72 bound to HsNMT1 so given the high degree of conservation of NMTs across species, we considered it preferable to use the *P.vivax* NMT as opposed to docking IMP72 into the crystal structure of IMP-917 in complex with HsNMT1. We used STRIFE to generate 250 elaborations for IMP-72, which we docked using the constrained docking functionality in GOLD, ^30^ ranking each compound by its ligand efficiency. Figure 4C shows the structure added to IMP-72 to create IMP-917 and several highly ranked elaborations proposed by STRIFE which appear to be capable of interacting with the Serine residue in the same way. Despite only generating a total of 250 compounds, some of the molecules proposed by STRIFE bear a striking resemblance with the trimethylpyrazole elaboration proposed by Mousnier et al. ^21^. A list of all unique elaborations generated by STRIFE can be found in the Supplementary Information (Figures S4-S8)

### Customisability

Whilst structure-aware generative models are increasingly being proposed, existing models incorporate such information through a single static structure, making them unable to account for the possibility that a side-chain may move to interact with a ligand. By utilising the flexible docking functionality in GOLD,^30^ STRIFE allows the user to explore design hypotheses where a specified side-chain moves; we illustrate how by considering a fragment-elaboration example from the literature.

O’Connell et al. ^22^ developed a small molecule inhibitor of tumour necrosis factor (TNF), a cytokine which has been shown to be a key factor in several autoimmune diseases, by making elaborations to a weakly binding fragment. The first elaboration allowed the formation of a hydrogen bond between the appended pyridyl group and the residue Y119^*A*^, which moved substantially in order to make the interaction, yielding a 2500-fold improvement in binding affinity (Figure 5a).

**Figure 5:**
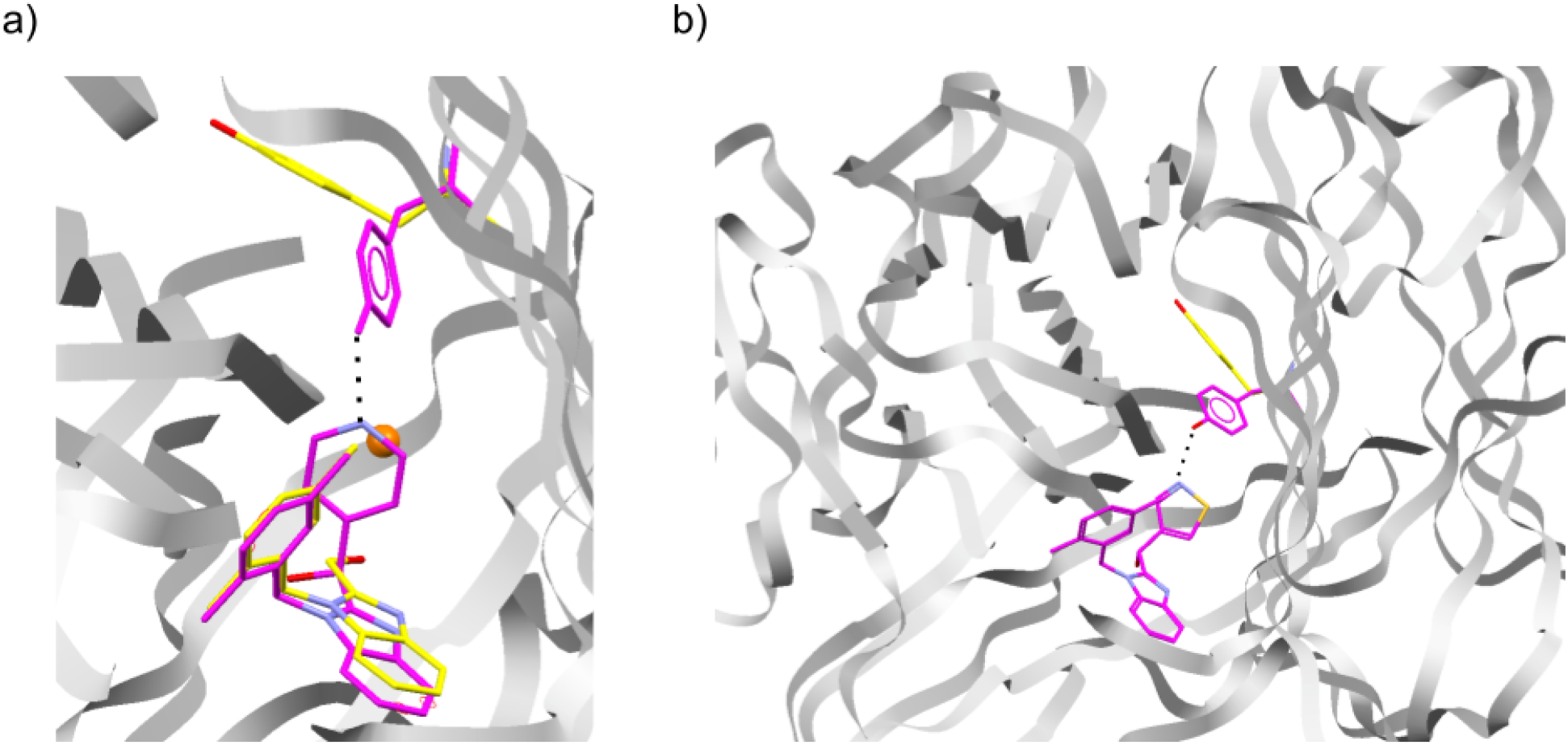
Visualisation of flexible docking using Hermes ^26^ a) Fragment (yellow carbons, PDB ID: 6OOY) with elaborated molecule (magenta carbons, PDB 6OOZ) reported by O’Connell et al. ^22^. The side chain of Y119^*A*^ moved substantially to form a hydrogen bond. The orange sphere represents a user-specified pharmacophoric point which we provided as input to STRIFE. b) An example of one of the molecules generated by STRIFE that appears to satisfy the specified design hypothesis. The molecule was docked into the fragment crystal structure (PDB ID: 6OOY, magenta side-chain is the predicted conformation) using the flexible docking functionality in GOLD, and supports the hypothesis that the side chain might move to accommodate the ligand.

The magnitude of the Y119^*A*^ side-chain movement presents a challenge for a generative model, as it would not be possible for a structure-aware model to predict that the sidechain would move, and if it was predicted by a chemist that the residue would be likely to move to form a hydrogen bond then it would not be possible to communicate such information to the generative model. Whilst STRIFE is unable to predict the movement of specific side-chains in advance, if a human expert has reason to believe a side-chain might move to accommodate a ligand, it is able to generate molecules which satisfy such a design hypothesis. This can be done by manually specifying a pharmacophoric point (see Methods, Customisability) such that a ligand pharmacophore placed at those coordinates would be able to interact with the residue side-chain if it were to move in the hypothesised fashion.

Under this set-up, STRIFE attempts to generate molecules with pharmacophores close to the user-specified pharmacophoric point and uses the flexible docking functionality in GOLD^30^ to dock the molecules whilst allowing the residue of interest to move freely; the user can then identify high scoring elaborations which were predicted to form the desired interaction with the protein.

To assess the ability of STRIFE to generate molecules which satisfied the design hypothesis specified by O’Connell et al. ^22^, we manually specified a pharmacophoric point (Figure 5a) and generated 250 elaborations using the same procedure as for our other experiments. To allow GOLD’s genetic algorithm to adequately explore the larger solution space created by side-chain flexibility, we generated 100 poses per molecule and used the highest scoring pose to calculate the corresponding ligand efficiency; further details of the flexible docking protocol can be found in the Supplementary Information.

STRIFE successfully recovered the highly potent pyridyl elaboration proposed by O’Connell et al. ^22^, whilst also proposing a wide range of structural analogues which appeared to be capable of making a similar hydrogen bond interaction with Y119^*A*^. In particular, the most common elaboration proposed by the model was a pyridine with a meta substitution pattern. In total, 49 of the 250 elaborations contain a pyridine substructure, whilst 125 elaborations included a hydrogen bond acceptor that was also part of an aromatic group. Elaborations comprising a six-membered aromatic ring with a hydrogen bond acceptor were not scored amongst the most ligand efficient using GOLD’s PLP scoring function, which generally rated pyrazole analogues or elaborations with hydrophobic groups more highly. However, consistent with the observed bound crystal structure, for both the ground-truth pyridyl elaboration and several highly-ranked elaborations which met the stated design hypothesis, the side-chain of Y119^*A*^ moved substantially to accommodate the proposed elaboration; an example is shown in Figure 5b and further details of the elaborations proposed by STRIFE can be found in the Supplementary Information (Figures S9, S10).

In summary, despite only making a small number of elaborations we were able to use the pharmacophoric information provided to make a range of plausible elaborations which satisfied the specified design hypothesis. In practice, predicting if and how a side chain may move is often extremely difficult but in such cases STRIFE can be used to assess the plausibility of such a movement and provide starting points for a fragment-to-lead campaign.

## Conclusion

We have proposed a model for fragment elaboration which derives meaningful information from the target into the generative process; unlike other generative models for fragment elaboration, STRIFE can incorporate target-specific information without using an existing active (although information from existing actives can easily be incorporated).

Currently, STRIFE uses information from FHMs which guide the placement of hydrogen bond acceptors and donors within the appended structure. Although hydrogen bonds between ligand and protein often lead to large improvements in binding affinity, they are by no means the only consideration when making elaborations to a fragment; the framework could easily be expanded to explicitly account for properties such as hydrophobicity and aromaticity, allowing a greater degree of control over the design process. A further limitation of the default implementation of STRIFE is that it does not seek to simultaneously satisfy multiple pharmacophoric points within a single elaboration, potentially curtailing its ability to generate highly efficient elaborations in some scenarios. However, fragment elaboration campaigns generally involve incrementally making small additions to the molecule and STRIFE provide the functionality to attempt to simultaneously satisfy multiple pharmacophoric points (whether FHM-derived or manually specified), should the user wish to.

Compared to existing structure-unaware models for fragment elaboration, the STRIFE algorithm carries a moderate up-front computational cost in calculating an FHM and identifying the set of quasi-actives (between 30-60 minutes on a desktop computer, in most cases). However, the most significant computational expense when generating a large number of elaborations is the docking of each generated molecule to estimate its ligand efficiency. As the quasi-actives only need to be identified once for a given fragment, the computational cost associated with STRIFE is therefore broadly comparable to other methods when generating large sets of molecules.

Although STRIFE is capable of being applied with minimal user input, one area which requires user specification is the choice of fragment and the associated exit vector. In practice, screening a fragment library may reveal dozens of weakly binding hits, yielding a large set of fragment-exit vector pairs to be explored; STRIFE could readily sample exhaustively from each fragment and exit vector, however a future avenue of research would be to develop a prioritisation scheme capable of identifying promising starting points for a fragmentto-lead campaign, to allow a more efficient allocation of resources.

An advantage of the representation of structural information that STRIFE extracts from the target is that it is extremely easy for a user to interpret. Whilst this is useful in allowing the user to understand why STRIFE generates the kinds of elaborations it does for a specific target, it also allows the user to easily specify their own design hypotheses. As such, we hope that STRIFE will be useful both in cases where a practitioner wishes to automatically generate a set of elaborations to a fragment bound to a novel target and in cases where they wish to rapidly enumerate a set of elaborations that conform to a specific design hypothesis and can be used as a basis for further designs.

## Code and Data Availability

STRIFE is available to download at https://github.com/oxpig/STRIFE. The default implementation of STRIFE is dependent on the commercial CSD Python API for calculating FHMs and carrying out constrained docking with GOLD. Users without access to the CSD Python API can still use STRIFE by manually specifying Pharmacophoric Points (see Methods, Customisability) and using alternative docking software.

SMILES strings of the molecules used to train the generative models and path length model can be accessed in the STRIFE github repository, as can the structures used for the large scale evaluation.

## Acknowledgements

T.E.H. is supported by funding from the Engineering and Physical Sciences Research Council (EPSRC), LifeArc, F. Hoffmann-La Roche AG, and UCB Pharma (Reference: EP/L016044/1). F.I. is supported by funding from EPSRC and Exscientia (Reference: EP/N509711/1). The authors would like to thank Mihaela Smilova, Ruben Sanchez-Garcia, Torsten Schindler, Lewis Vidler, Will Pitt and Garrett M. Morris for helpful discussions.

