## Supplementary Info for "Incorporating Target-Specific Pharmacophoric Information Into Deep Generative Models For Fragment Elaboration"

### Pharmacophoric Profiles

To make the notion of pharmacophoric profiles more concrete, we provide examples of pharmacophoric profiles which could be provided to the generative model and examples of elaborations which conform to the specified profile and examples of molecules which do not conform to the profile.

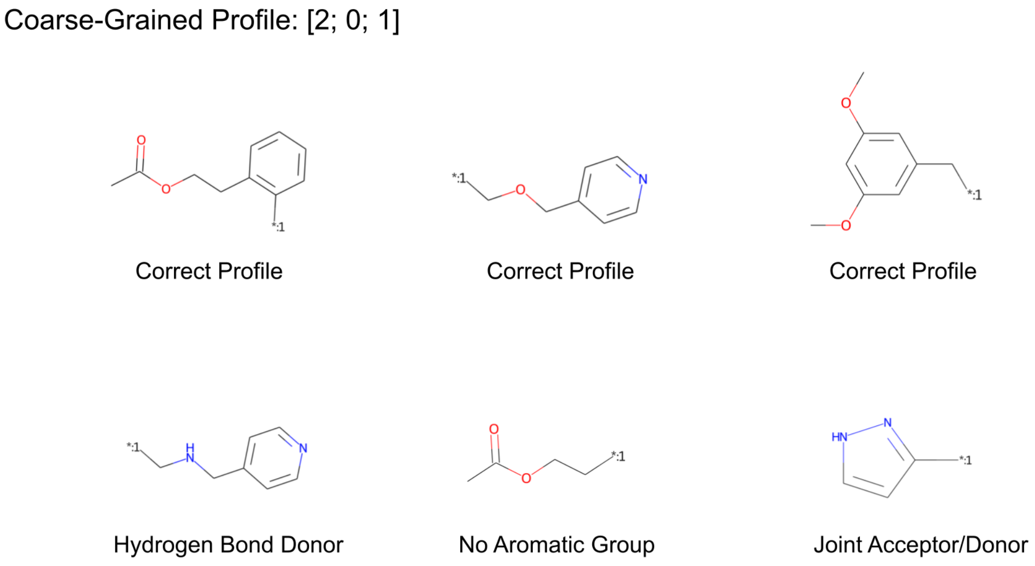


Figure S1: The coarse-grained profile is specified as [# Hydrogen Bond Acceptors; # Hydrogen Bond Donors; # Aromatic Groups]. The elaborations in the first row all have two Hydrogen Bond Acceptors, no Hydrogen Bond Donors and an Aromatic group, whilst the elaborations in the second row fail to conform to the specified profile.

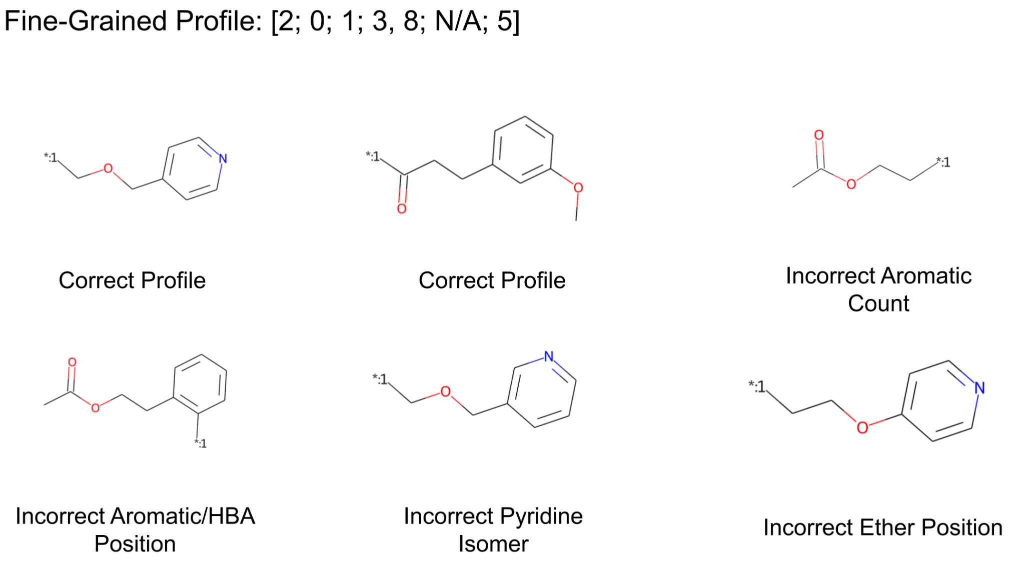


Figure S2: The fine-grained profile is specified as [# Hydrogen Bond Acceptors; # Hydrogen Bond Donors; # Aromatic Groups; Path Distance of Acceptors; Path Distance of Donors; Path Distance to first Aromatic Atom]. Path distances are inclusive of both the exit vector and the pharmacophore. The first two in the top row satisfy the specified fine-grained profile, whereas the third elaboration does not include the correct number of pharmacophores. Each elaboration in the bottom row fails to place the desired pharmacophores in the correct locations.

#### Filtering in the Exploration phase

We apply two filtering stages in the Exploration phase of the STRIFE algorithm:

- Molecules are filtered out if they do not have the same pharmacophoric counts as provided in the coarse-grained pharmacophoric profile.
- In addition molecules must pass the 2D filters described in (Methods, Evaluation Metrics).

### Examples in the CASF Test Set

| PDB ID | Fragment SMILES String |
|---|---|
| 1pxn | CNc1nc(C)c(s1)-c1ccnc(n1)N[*:1] |
| 1q8t | CC([NH3+])C1CCC(CC1)C(=O)N[*:1] |
| 1uto | c1ccc(cc1)[*:1] |
| 1ydt | O=S(=O)(NCC[NH2+]C[*:1])c1cccc2cnccc12 |
| 1ydt | O=S(=O)(NCC[NH2+]CC=C[*:1])c1cccc2cnccc12 |
| 1z95 | CC(O)(CS(=O)(=O)c1ccc(F)cc1)C(=O)Nc1ccc(C#N)c(c1)[*:1] |
| 2br1 | COc1ccc(cc1)-c1oc2ncnc(c2c1-c1ccc(cc1)OC)[*:1] |
| 2br1 | COc1ccc(cc1)-c1oc2ncnc(N[*:1])c2c1-c1ccc(cc1)OC |
| 2br1 | COc1ccc(cc1)-c1c(oc2ncnc(NCCO)c12)[*:1] |
| 2brb | c1ccc(cc1)-c1oc2ncnc(N[*:1])c2c1-c1ccccc1 |
| 2brb | c1ccc(cc1)-c1oc2ncnc(c2c1-c1ccccc1)[*:1] |
| 2c3i | CC(=O)c1cccc(c1)-c1cnc2ccc(nn12)NC[*:1] |
| 2c3i | CC(=O)c1cccc(c1)-c1cnc2ccc(nn12)N[*:1] |
| 2c3i | CC(=O)c1cccc(c1)-c1cnc2ccc(nn12)[*:1] |
| 2cet | OCC1C(O)C(O)C(O)c2nc(cn21)C[*:1] |
| 2cet | OCC1C(O)C(O)C(O)c2nc(cn21)CC[*:1] |
| 2fvd | CS(=O)(=O)N1CCC(CC1)Nc1ncc(C(=O)[*:1])c(N)n1 |
| 2fvd | COc1ccc(F)c(F)c1C(=O)c1cnc(nc1N)N[*:1] |
| 2p15 | CC12CCC3c4ccc(O)cc4CCC3C1CCC2(O)C=C[*:1] |
| 2p15 | CC12CCC3c4ccc(O)cc4CCC3C1CCC2(O)/C=C/c1ccccc1[*:1] |
| 2pog | Oc1cccc2OC(C3CCCC3c12)[*:1] |
| 2qe4 | COCc1cc(O)cc2c1OC(C1CCCC21)[*:1] |
| 2w66 | OCC1[NH2+]CC(O)C(C(O)C1O)[*:1] |
| 2wbg | OC1C(O)C(O)N2/C(=N/CCC[*:1])OCC2C1O |

| | |
|---|---|
| 2wbg | OC1C(O)C(O)N2/C(=N/C[*:1])OCC2C1O |
| 2wbg | OC1C(O)C(O)N2/C(=N/CCCC[*:1])OCC2C1O |
| 2wbg | OC1C(O)C(O)N2/C(=N/CC[*:1])OCC2C1O |
| 2wbg | OC1C(O)C(O)N2/C(=N/CCCCC[*:1])OCC2C1O |
| 2wn9 | COc1cc(O)ccc1/C=C1\CCCN=C1[*:1] |
| 2xbv | O=C(Nc1ccc(cc1F)-n1ccccc1=O)C1C[NH+](CC(F)F)CC1[*:1] |
| 2xbv | O=C(Nc1ccc(cc1F)-n1ccccc1=O)C1C[NH+](CC(F)F)CC1C(=O)N[*:1] |
| 2xbv | O=C(Nc1ccc(Cl)cn1)C1C[NH+](CC(F)F)CC1C(=O)Nc1ccc(cc1F)[*:1] |
| 2xj7 | OC1CC[NH+]2CC(C(O)C(O)C12)[*:1] |
| 2yki | O=C(NC1c2ccccc2-c2c(cccc21)-c1nc2ccncc2[nH]1)[*:1] |
| 2yki | O=C(NC1c2ccccc2-c2c1cccc2[*:1])c1ccnc2[nH]ccc12 |
| 2ymd | Oc1ccc2[nH]cc(c2c1)[*:1] |
| 2zb1 | Cc1nnc(o1)-c1ccc(C)c(c1)-c1ccc(cc1)C(=O)N[*:1] |
| 2zb1 | Cc1nnc(o1)-c1ccc(C)c(c1)-c1ccc(cc1)C(=O)NC[*:1] |
| 2zb1 | Cc1nnc(o1)-c1ccc(C)c(c1)-c1ccc(cc1)[*:1] |
| 2zb1 | Cc1ccc(cc1-c1ccc(cc1)C(=O)NCC1CC1)[*:1] |
| 3acw | OC1(C[NH+]2CCC1CC2)c1ccc(cc1)[*:1] |
| 3aru | Cn1cnc2c1c(=O)n(CC[*:1])c(=O)n2C |
| 3aru | Cn1cnc2c1c(=O)n(C[*:1])c(=O)n2C |
| 3aru | Cn1cnc2c1c(=O)n(CCCC[*:1])c(=O)n2C |
| 3aru | Cn1cnc2c1c(=O)n(CCC[*:1])c(=O)n2C |
| 3ary | c1cc2oc(cc2cc1[*:1])C1=NCCN1 |
| 3b1m | COc1cc(O)c2c(OC3=CC(O)=C(C(C)=O)C(=O)C32C)c1C(=O)NC[*:1] |
| 3b5r | CC(O)(CO[*:1])C(=O)Nc1ccc(C#N)c(c1)C(F)(F)F |
| 3b65 | CC(O)(CO[*:1])C(=O)Nc1ccc(C#N)c(I)c1 |
| 3b68 | CC(O)(CO[*:1])C(=O)Nc1ccc(c(c1)C(F)(F)F)[N+](=O)[O-] |
| 3b68 | CC(O)(COc1ccc(cc1)[*:1])C(=O)Nc1ccc(c(c1)C(F)(F)F)[N+](=O)[O-] |

| | |
|---|---|
| 3b68 | CC(=O)Nc1ccc(cc1)OCC(C)(O)C(=O)Nc1ccc(c(c1)[*:1])[N+](=O)[O-] |
| 3coy | CC(C)(C)C([NH3+])C(=O)NS(=O)(=O)OCC1OC(C(O)C1O)[*:1] |
| 3coy | Nc1ncnc2c1ncn2C1OC(COS(=O)(=O)NC(=O)C([NH3+])[*:1])C(O)C1O |
| 3coz | Nc1ncnc2c1ncn2C1OC(COS(=O)(=O)NC(=O)C([NH3+])[*:1])C(O)C1O |
| 3coz | CC(C)C([NH3+])C(=O)NS(=O)(=O)OCC1OC(C(O)C1O)[*:1] |
| 3e92 | Cc1cc(ccc1-c1cc(ccc1C)C(=O)NC1CC1)[*:1] |
| 3e92 | Cc1nnc(o1)-c1ccc(c(C)c1)-c1cc(ccc1C)[*:1] |
| 3fur | O=S(=O)(Nc1cc(Cl)c(Oc2cnc3ccccc3c2)c(Cl)c1)[*:1] |
| 3fur | O=S(=O)(Nc1cc(Cl)c(O[*:1])c(Cl)c1)c1ccc(Cl)cc1Cl |
| 3g0w | Cc1c(ccc(C#N)c1Cl)/N=C1\OC(C2C(O)CCN12)[*:1] |
| 3g2n | O=C(NC1OC(CO)C(O)C(O)C1O)[*:1] |
| 3lka | COc1ccc(cc1)[*:1] |
| 3nx7 | COc1ccc(cc1)S(=O)(=O)N(CCO)[*:1] |
| 3nx7 | COc1ccc(cc1)S(=O)(=O)N(CCO)C[*:1] |
| 3qgy | c1ccc(cc1)-c1cc2c(ccc3cnc(nc32)Nc2cccc(c2)[*:1])s1 |
| 3qgy | c1ccc(cc1)-c1cc2c(ccc3cnc(nc32)N[*:1])s1 |
| 3rsx | Nc1ccc2cc(ccc2n1)[*:1] |
| 3u8k | c1ncc(cc1N1CCC[NH2+]CC1)[*:1] |
| 3u8n | Brc1ncc(cc1[*:1])N1CCC[NH2+]CC1 |
| 4dli | Nc1ccc2c(nc(nc2c1)-c1ccccc1)[*:1] |
| 4dli | Nc1ccc2c(nc(nc2c1)[*:1])NC1CC1 |
| 4dli | Nc1ccc2c(nc(nc2c1)-c1ccccc1)N[*:1] |
| 4e6q | c1cc2c(ncc3ncn(c32)C2CC[NH+](CC2)C[*:1])[nH]1 |
| 4ea2 | C/C(=C/C[NH2+]CC[NH2+]C1C2CC3CC(C2)CC1C3)C[*:1] |
| 4eky | O=C1NC(=O)N(CC1=C=C[*:1])C1OC(CO)C(O)C(O)C1O |
| 4eor | c1nc2c(nc(nc2[nH]1)Nc1ccc(cc1)[*:1])OCC1CCCCC1 |
| 4eor | NS(=O)(=O)c1ccc(cc1)Nc1nc(c2nc[nH]c2n1)[*:1] |

| | |
|---|---|
| 4eor | NS(=O)(=O)c1ccc(cc1)Nc1nc(OC[*:1])c2nc[nH]c2n1 |
| 4eor | NS(=O)(=O)c1ccc(cc1)Nc1nc(O[*:1])c2nc[nH]c2n1 |
| 4f3c | Nc1ncnc2c(c[nH]c12)C[NH+]1CC(O)C(CSC[*:1])C1 |
| 4f9w | c1ccc2cc(ccc2c1)-c1nnc(cc1-c1ccncc1)[*:1] |
| 4f9w | CN(C)c1cc(c(nn1)-c1ccc2ccccc2c1)[*:1] |
| 4ih7 | O=C1N=CC=CC1c1cccc(c1)[*:1] |
| 4ivb | N#CC1CCC(CC1)n1c(nc2cnc3[nH]ccc3c21)[*:1] |
| 4ivd | N#CCCC1CCC(CC1)n1c(nc2cnc3[nH]ccc3c21)[*:1] |
| 4ivd | CC(O)c1nc2cnc3[nH]ccc3c2n1C1CCC(CC1)[*:1] |
| 4ivd | CC(O)c1nc2cnc3[nH]ccc3c2n1C1CCC(CC1)C[*:1] |
| 4lzs | CCc1c([nH]c(C)c1[*:1])C(=O)NC |
| 4lzs | CCc1c([nH]c(C)c1C(C)=O)[*:1] |
| 4m0y | NC(=O)Nc1cn(nc1[*:1])-c1cccc2ccccc12 |
| 4m0z | COc1ccc2cccc(c2c1)-n1cc(c(n1)C(N)=O)[*:1] |
| 4m0z | COc1ccc2cccc(c2c1)-n1cc(NC(N)=O)c(n1)[*:1] |
| 4mgd | Oc1ccc(cc1)C(c1ccc(O)cc1)[*:1] |
| 4qac | Nc1nc(cc(n1)[*:1])-c1ccc(cc1)C(F)(F)F |
| 4twp | CNC(=O)c1ccccc1Sc1ccc2c(C=C[*:1])[nH]nc2c1 |
| 4twp | C(=C/c1[nH]nc2cc(ccc12)Sc1ccccc1[*:1])\c1ccccn1 |
| 4twp | CNC(=O)c1ccccc1Sc1ccc2c([nH]nc2c1)[*:1] |
| 5c2h | Cc1c(Cl)nc(nc1NC[*:1])OCCCc1ccc2ccccc2n1 |
| 5c2h | Cc1c(Cl)nc(nc1N[*:1])OCCCc1ccc2ccccc2n1 |
| 5c2h | Cc1nc(C)c(CNc2nc(nc(Cl)c2C)OCCC[*:1])s1 |

Table S1: Examples used for the large scale evaluation.

### Additional results

Table S2: Comparison of results obtained by CReM, on different sets of examples.

| Metric | All Examples | > 50 Elaborations | 250 Elaborations |
|---|---|---|---|
| Valid | **100%** | **100%** | **100%** |
| Unique | N/A | N/A | N/A |
| Novel | N/A | N/A | N/A |
| Pass 2D filters | **67.44%** | 66.94% | 66.06% |
| $\Delta SLE_{20}$ | -2.015 | -0.037 | **-0.029** |
| $\Delta SLE_{50}$ | -2.443 | -0.536 | **-0.489** |
| $\Delta SLE_{100}$ | -2.812 | -1.024 | **-0.992** |

When including examples where CReM proposed fewer than 50 examples into the calculation of summary statistics for CReM ("All Examples"), the method's performance degrades substantially. This is primarily due to instability in the scaling procedure for small sample sizes, where the ground truth ligand efficiency was standardised to be extremely large.

#### Effect of changing $\alpha$ in $\Delta SLE_\alpha$

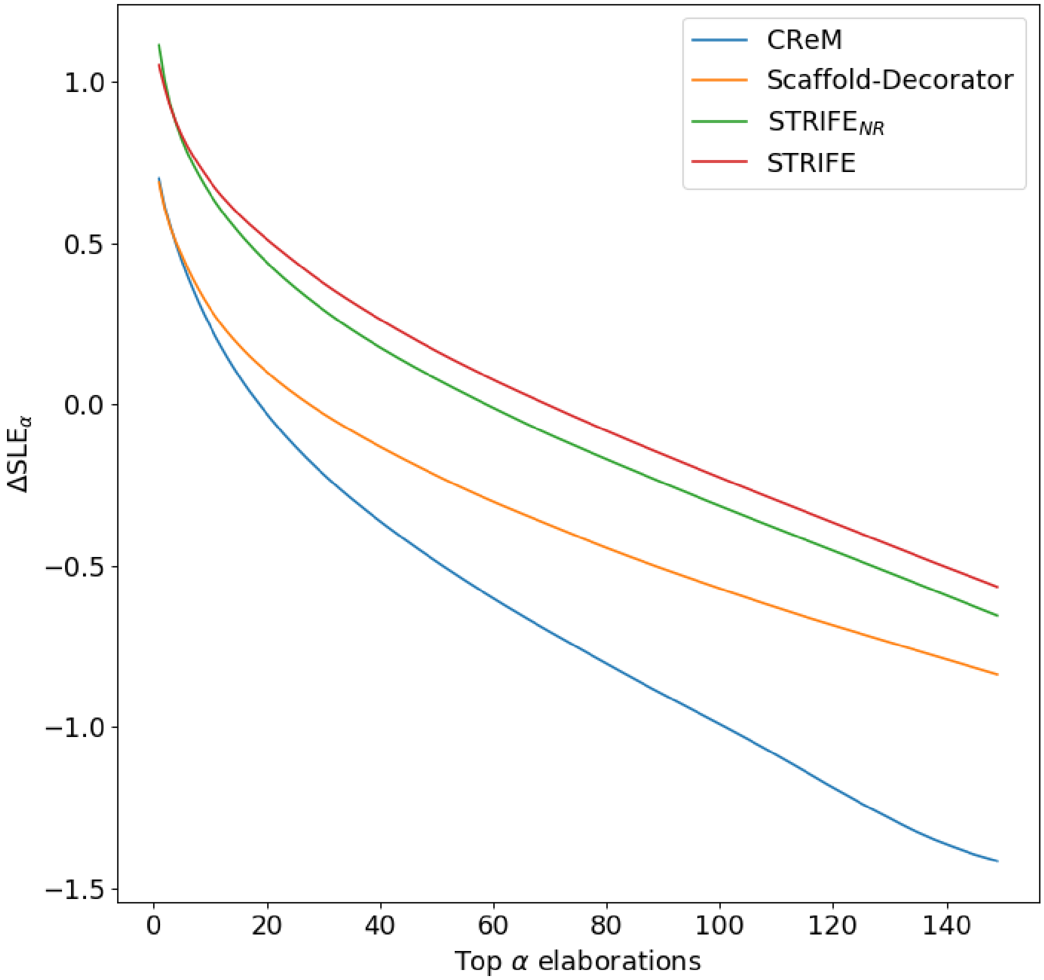


Figure S3: Effect of changing the proportion of elaborations made by each method on the Standardised Ligand Efficiency Improvement ($\Delta SLE_\alpha$). Unsurprisingly, as more elaborations are included in the calculation, the average ligand efficiency is degraded compared to the ground truth.

### Generated Molecules

We present all unique molecules generated in both case studies, ranked by their ligand efficiency.

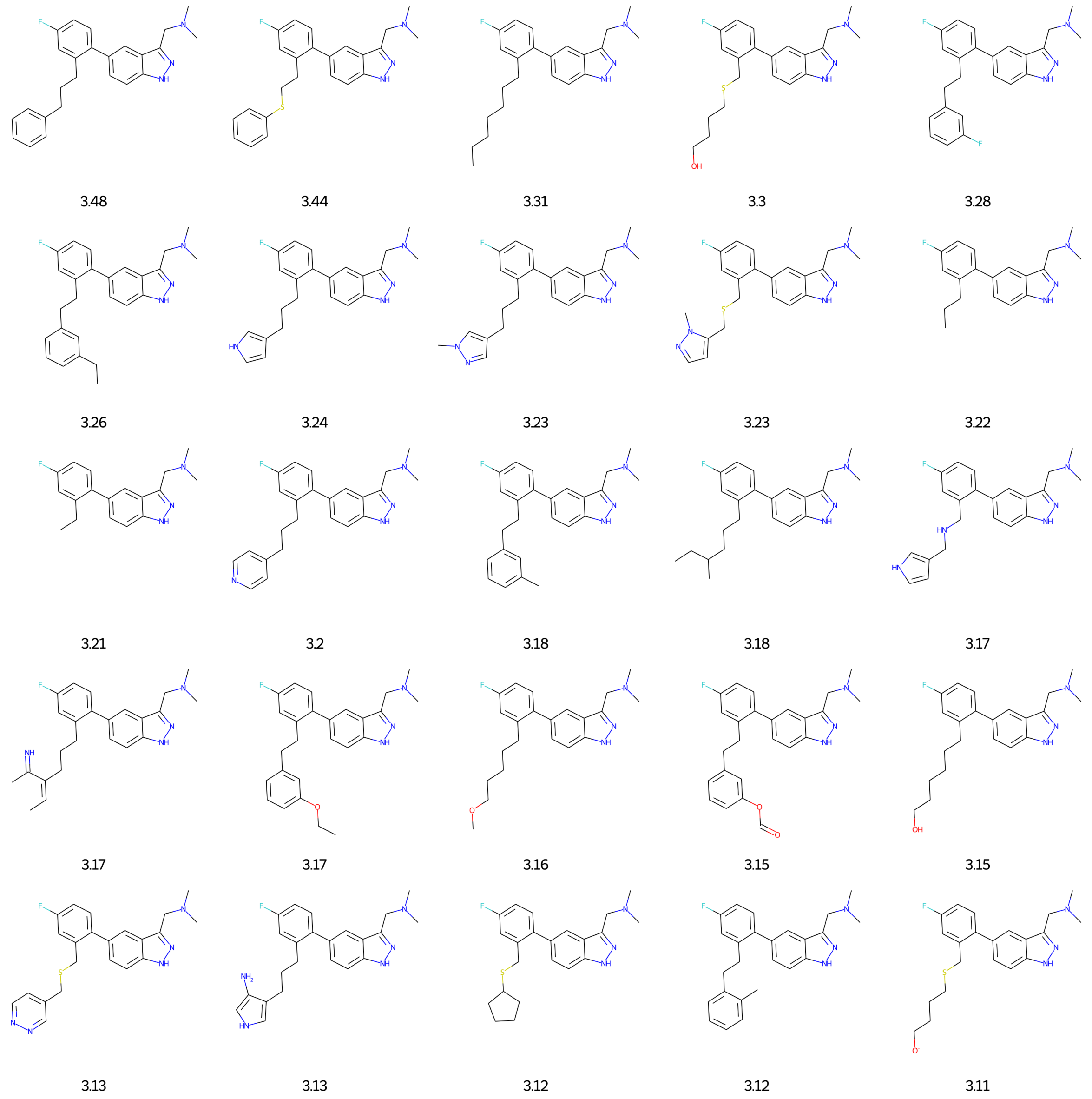


Figure S4: Unique molecules generated on first case study with associated ligand efficiency: Rank 1-25

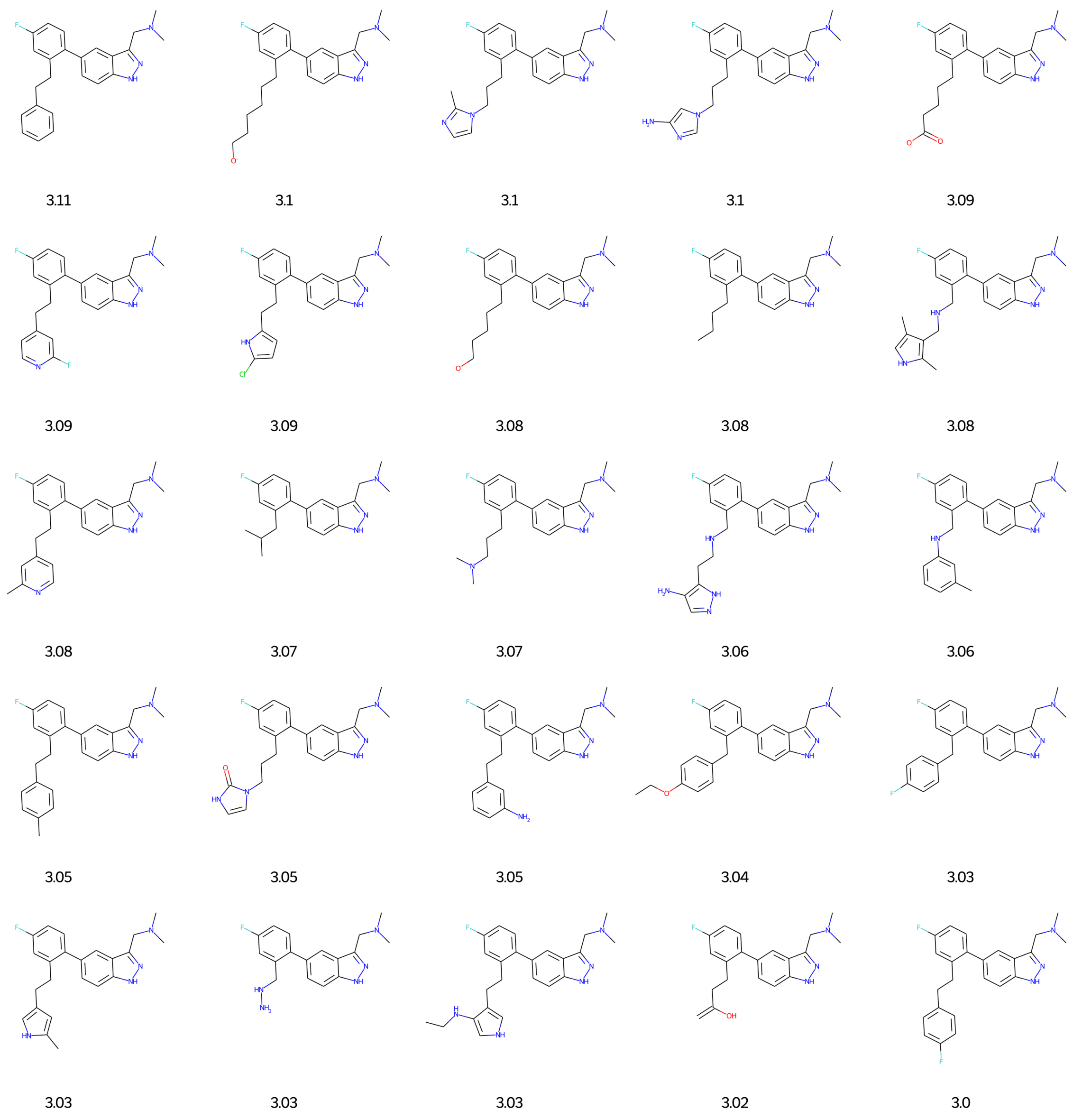


Figure S5: Unique molecules generated on first case study with associated ligand efficiency: Rank 26-50

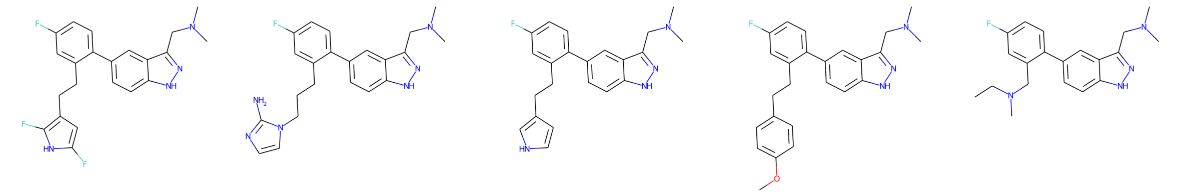

3.0 | 2.99 | 2.98 | 2.98 | 2.97

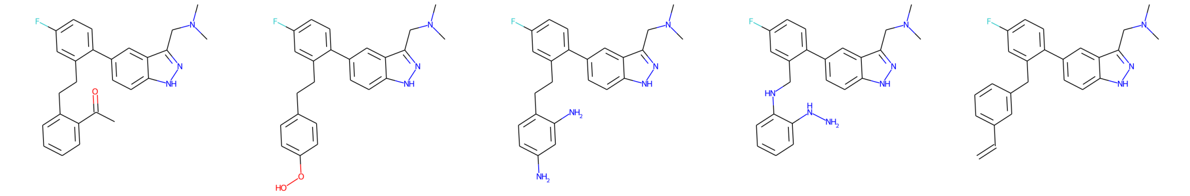

2.96 | 2.94 | 2.91 | 2.9 | 2.9

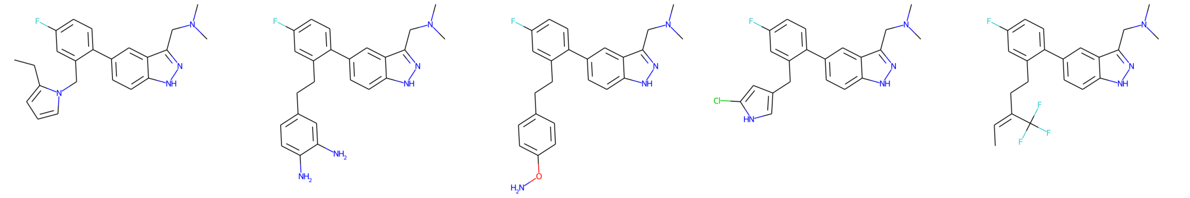

2.88 | 2.88 | 2.87 | 2.87 | 2.87

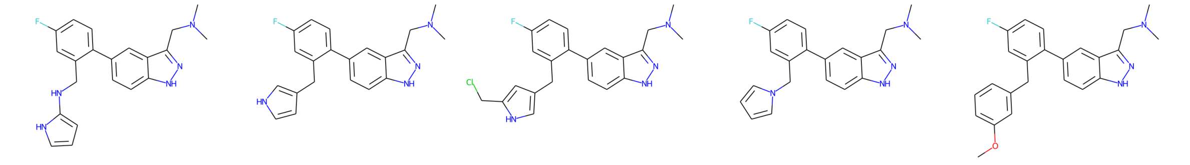

2.86 | 2.85 | 2.84 | 2.83 | 2.83

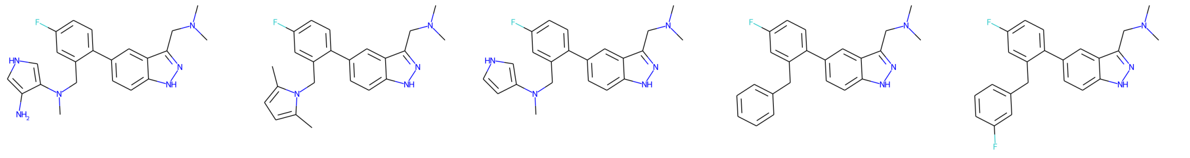

2.83 | 2.82 | 2.82 | 2.8 | 2.8

Figure S6: Unique molecules generated on first case study with associated ligand efficiency: Rank 51-75

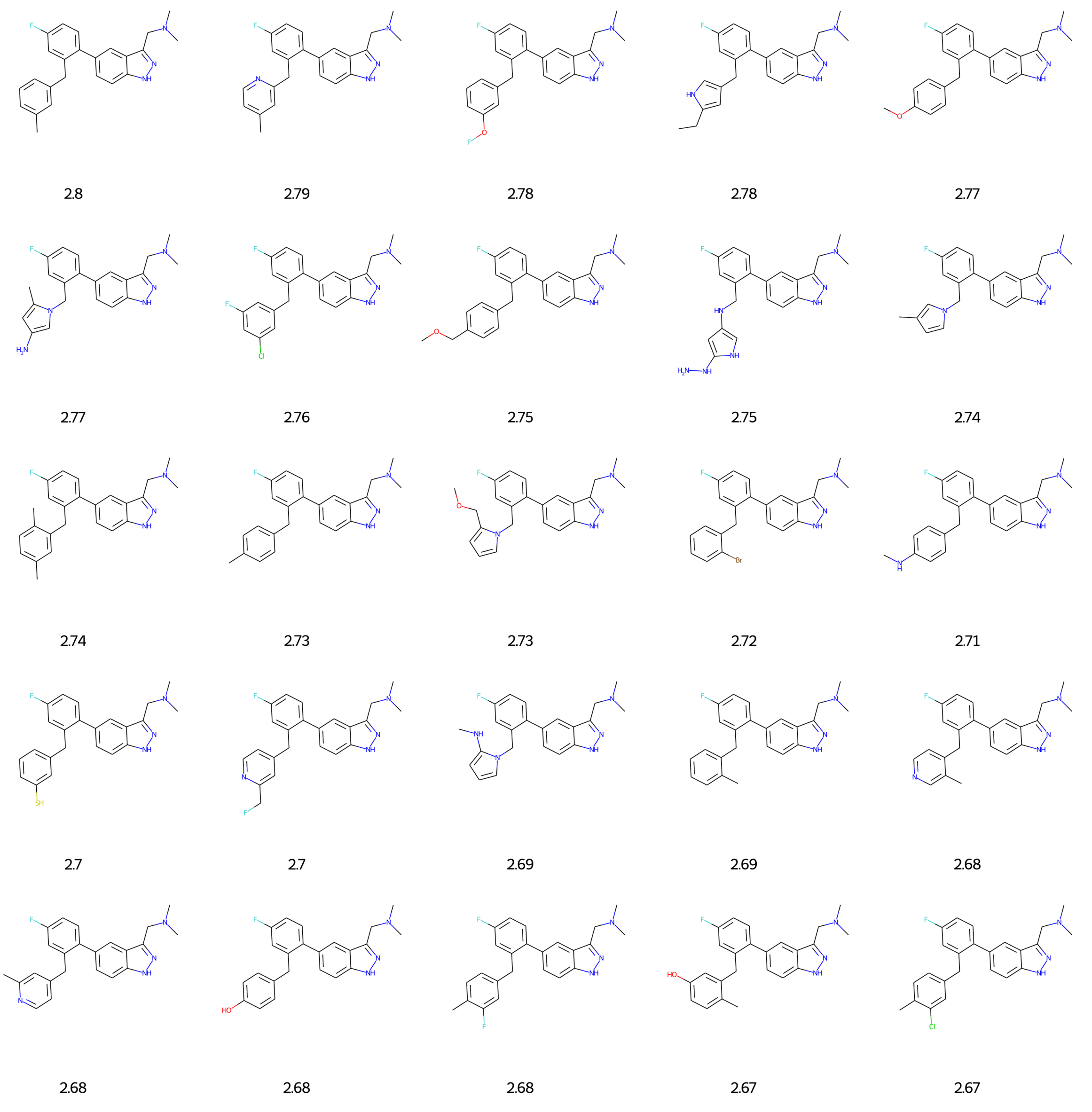


Figure S7: Unique molecules generated on first case study with associated ligand efficiency: Rank 76-100

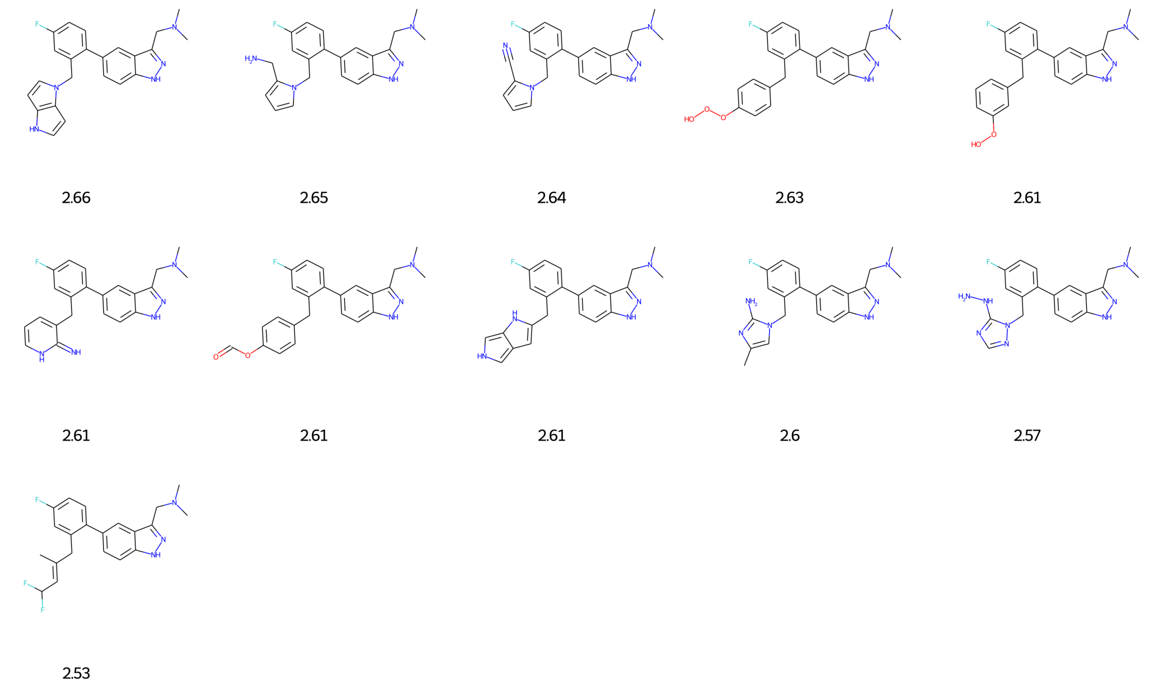


Figure S8: Unique molecules generated on first case study with associated ligand efficiency: Rank 101-111

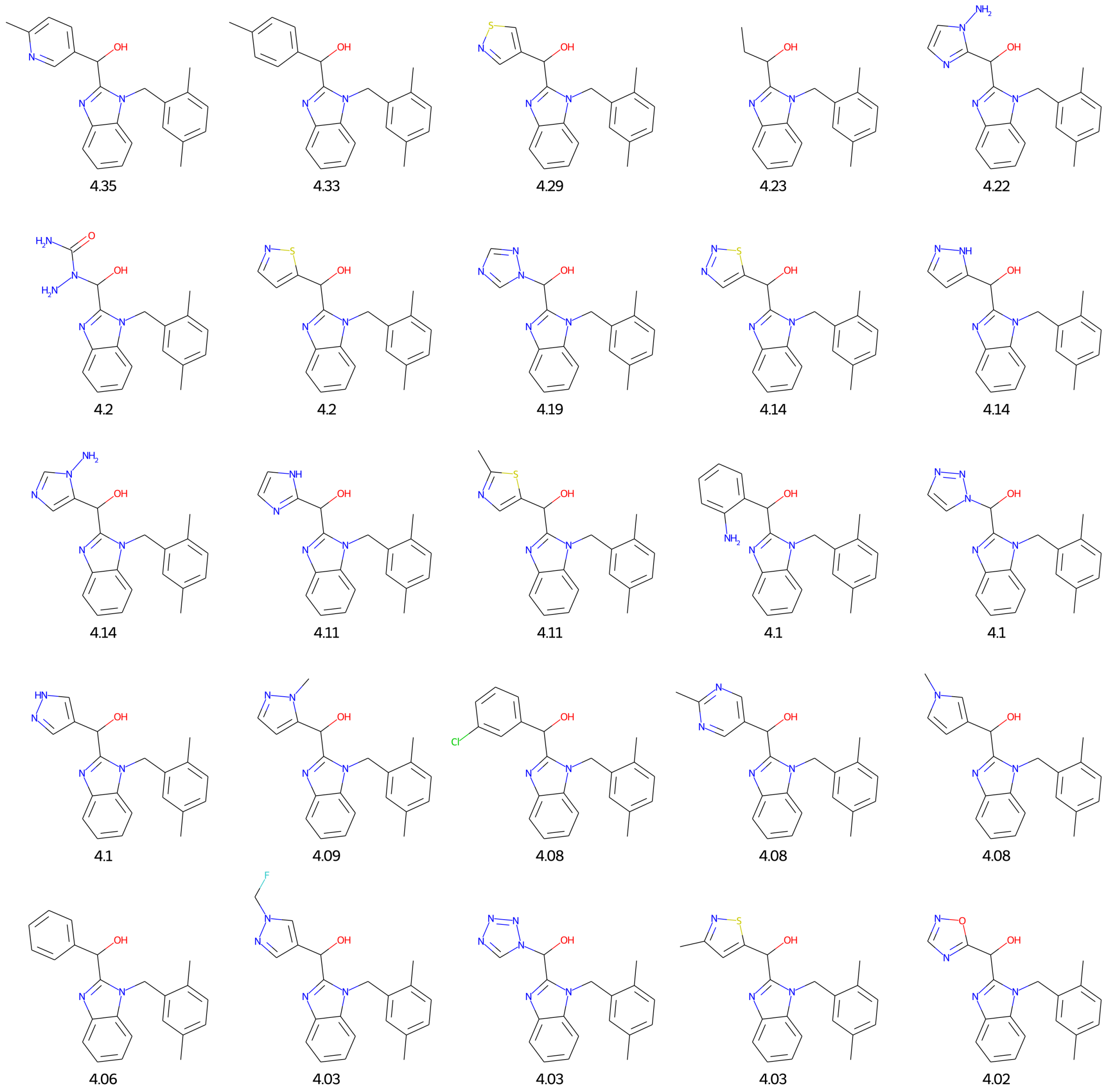


Figure S9: Unique molecules generated on second case study with associated ligand efficiency: Rank 1-25

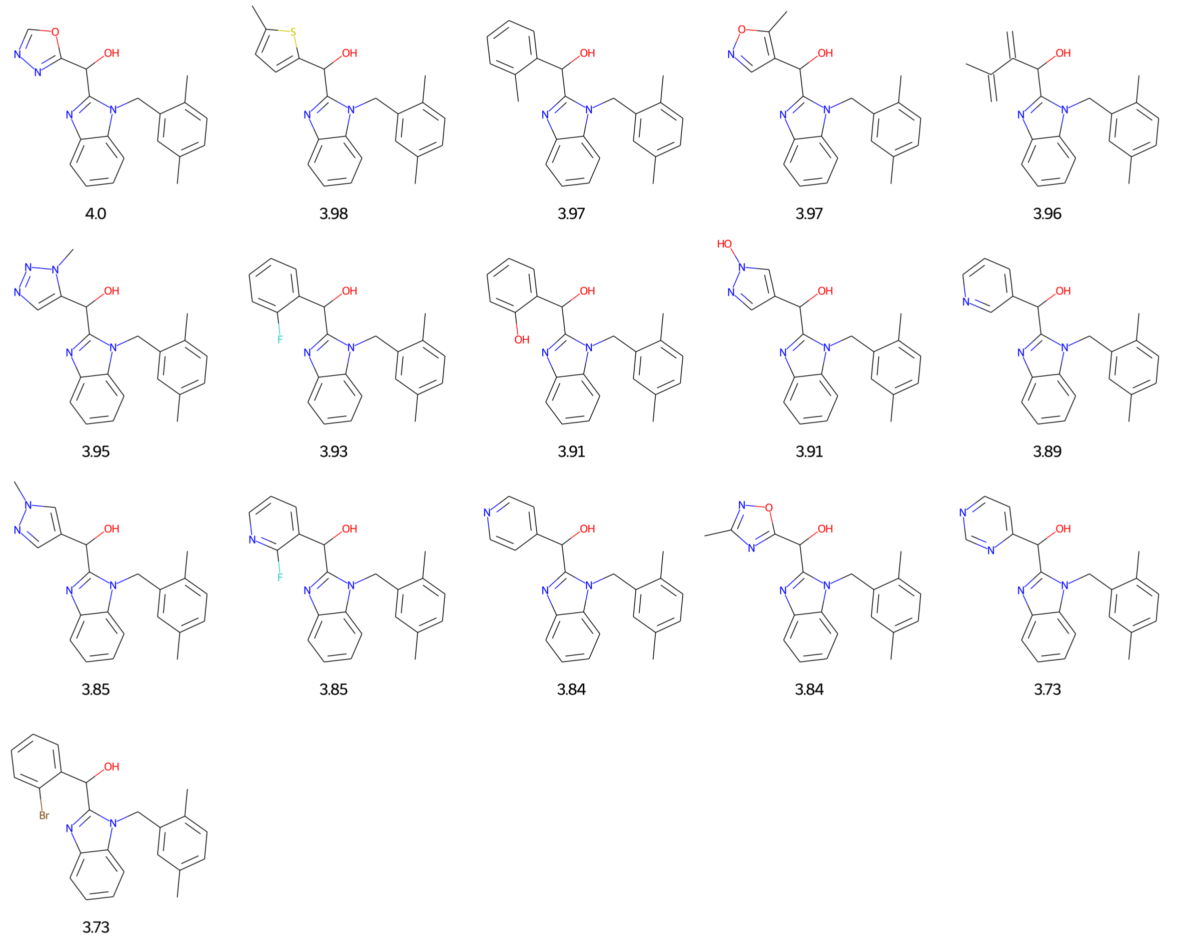


Figure S10: Unique molecules generated on second case study with associated ligand efficiency: Rank 26-44

### GOLD Flexible Docking Configuration File

The following configuration file was used to carry out flexible docking in GOLD..[1]

```
GOLD CONFIGURATION FILE


AUTOMATIC SETTINGS
autoscale = 0.1


POPULATION
popsiz = auto
select_pressure = auto
n_islands = auto
maxops = auto
niche_siz = auto


GENETIC OPERATORS
pt_crosswt = auto
allele_mutatewt = auto
migratewt = auto


FLOOD FILL
radius = 10
origin = 0 0 0
do_cavity = 1
floodfill_atom_no = 0
cavity_file = <path/to/cavity/file>.mol2
floodfill_center = cavity_from_ligand 10 atoms
```

DATA FILES

```
ligand_data_file <path/to/ligands>.sdf 100
param_file = DEFAULT
set_ligand_atom_types = 1
set_protein_atom_types = 0
directory = <output_directory>
tordist_file = DEFAULT
make_subdirs = 0
save_lone_pairs = 1
fit_points_file = fit_pts.mol2
read_fitpts = 0
```

FLAGS

```
internal_ligand_h_bonds = 0
flip_free_corners = 0
match_ring_templates = 0
flip_amide_bonds = 0
flip_planar_n = 1 flip_ring_NRR flip_ring_NHR
flip_pyramidal_n = 0
rotate_carboxylic_oh = flip
use_tordist = 1
postprocess_bonds = 1
rotatable_bond_override_file = DEFAULT
solvate_all = 1
```

TERMINATION

```
early_termination = 0
n_top_solutions = 3
rms_tolerance = 1.5

  CONSTRAINTS
force_constraints = 0
constraint scaffold <path/to/scaffold>.mol2 5.0000

  COVALENT BONDING
covalent = 0

  SAVE OPTIONS
save_score_in_file = 1 comments
save_protein_torsions = 1
concatenated_output = <output_directory>/docked_ligands.sdf
output_file_format = MACCS

  FITNESS FUNCTION SETTINGS
initial_virtual_pt_match_max = 3
relative_ligand_energy = 1
gold_fitfunc_path = plp
score_param_file = DEFAULT

  PROTEIN DATA
protein_datafile = <path/to/protein>/6ooy_protein.pdb

rotamer_lib
```

```
  name TYR119
 chi1 1586 1585 1587 1586
 chi2 1585 1586 1589 1590
 chi3 1586 1589 1590 1591
 rotamer 0 (60) 0 (180) 0 (180)
end_rotamer_lib

rotamer_lib
  name LEU120
 chi1 1607 1606 1608 1607
 chi2 1606 1607 1610 1611
 chi3 1607 1610 1611 1613
 rotamer 0 (60) 0 (180) 0 (180)
end_rotamer_lib
```
